# Viral channelrhodopsins: calcium-dependent Na^+^/K^+^ selective light-gated channels

**DOI:** 10.1101/2020.02.14.949966

**Authors:** D. Zabelskii, A. Alekseev, K. Kovalev, A.-S. Oliviera, T. Balandin, D. Soloviov, D. Bratanov, D. Volkov, S. Vaganova, R. Astashkin, I. Chizhov, N. Yutin, M. Rulev, A. Popov, T. Rokitskaya, Y. Antonenko, R. Rosselli, F. Rodriguez-Valera, G. Armeev, K. Shaitan, G. Bueldt, M. Vivaudou, M. Kirpichnikov, E. Koonin, E. Bamberg, V. Gordeliy

**Affiliations:** Institute of Biological Information Processing (IBI-7: Structural Biochemistry), Forschungszentrum Jülich, Jülich, Germany; JuStruct: Jülich Center for Structural Biology, Forschungszentrum Jülich, Jülich, Germany; Research Center for Molecular Mechanisms of Aging and Age-related Diseases, Moscow Institute of Physics and Technology, Dolgoprudny, Russia; Institute of Crystallography, University of Aachen (RWTH), Aachen, Germany; Institut de Biologie Structurale J.-P. Ebel, Université Grenoble Alpes-CEA-CNRS, Grenoble, France; Joint Institute for Nuclear Research, Dubna, Russia; Institute for Safety Problems of Nuclear Power Plants, NAS of Ukraine, Kyiv, 03680, Ukraine; Institute for Biophysical Chemistry, Hannover Medical School, Hannover, Germany; National Center for Biotechnology Information, National Library of Medicine, National Institutes of Health, Bethesda, MD, United States; European Synchrotron Radiation Facility Grenoble, France; Belozersky Institute of Physico-Chemical Biology, Lomonosov Moscow State University, Moscow, Russia; Evolutionary Genomics Group, Departamento de Producción Vegetal y Microbiología, Universidad Miguel Hernández, San Juan de Alicante, Spain; School of Biology, Lomonosov Moscow State University, Moscow 119991, Russia; Laboratories of Excellence, Ion Channel Science and Therapeutics, 06560 Valbonne, France; Max Planck Institute of Biophysics, Frankfurt am Main, Germany

**Author notes:** These authors contributed equally to this work (D. Zabelskii, A. Alekseev, K. Kovalev). Corresponding author. (V.G.).

## Abstract

Phytoplankton is the base of the marine food chain, oxygen, carbon cycle playing a global role in climate and ecology. Nucleocytoplasmic Large DNA Viruses regulating the dynamics of phytoplankton comprise genes of rhodopsins of two distinct families. We present a function-structure characterization of two homologous proteins representatives of family 1 of viral rhodopsins, OLPVR1 and VirChR1. VirChR1 is a highly selective, Ca^2+^-dependent, Na^+^/K^+^- conducting channel and, in contrast to known cation channelrhodopsins (ChRs), is impermeable to Ca^2+^ ions. In human neuroblastoma cells, upon illumination, VirChR1 depolarizes the cell membrane to a level sufficient to fire neurons. It suggests its unique optogenetic potential. 1.4 Å resolution structure of OLPVR1 reveals their remarkable difference from the known channelrhodopsins and a unique ion-conducting pathway. The data suggest that viral channelrhodopsins mediate phototaxis of algae enhancing the host anabolic processes to support virus reproduction, and therefore, their key role in global phytoplankton dynamics.

## Introduction

Microbial and animal rhodopsins (type-1 and 2 rhodopsins, respectively) comprise a superfamily of heptahelical (7-TM) transmembrane proteins covalently linked to a retinal chromophore (Ernst et al., 2014; Gushchin and Gordeliy, 2018). Type-1 rhodopsins are the most abundant light-harvesting proteins that have diverse functions, such as ion pumping, ion channeling, sensory and enzymatic activities (Gordeliy et al., 2002; Govorunova et al., 2015; Gushchin et al., 2013, 2015; Kato et al., 2015; Kim et al., 2018; Mukherjee et al., 2019; Shevchenko et al., 2017; Volkov et al., 2017). The discovery, in 2000, of the light-driven pump proteorhodopsin (PR) in marine microbes triggered extensive search of metagenomes for light-activated proteins (Béjà et al., 2000). As a result, about 10,000 rhodopsin genes have been identified in archaea, bacteria, unicellular eukaryotes and viruses although the biological functions of most of these proteins remains elusive. The study of microbial rhodopsins culminated in the development of optogenetics, the revolutionary method for controlling cell behavior in vivo using light-gated channels and light-driven pumps (Deisseroth, 2015). Currently, major efforts are being undertaken to identify rhodopsins with novel functions and properties that could be harnessed to enhance optogenetic applications (Berndt and Deisseroth, 2015; Berndt et al., 2014; Govorunova et al., 2015; Kato et al., 2018; Kim et al., 2018; Sineshchekov et al., 2017a).

In 2012, bioinformatic analysis of proteins encoded by Nucleocytoplasmic Large DNA Viruses (NCLDV) resulted in the identification of highly diverged PR homologs (hereafter, viral rhodopsins) in Organic Lake Phycodnavirus and Phaeocystis globosa viruses which belong to the extended Mimiviridae family (Yutin and Koonin, 2012). Phylogenetic analysis shows that, within the rhodopsin superfamily, viral rhodopsins form a monophyletic group that consists of two distinct branches, VR1 and VR2 (López et al., 2017). Recently, a DTS-rhodopsin from VR1 group was reported to pump protons when expressed in E.coli plasma membrane (Needham et al., 2019). In contrast to VR1 group, the OLPVRII protein from VR2 group was shown to have pentameric organization that resembles ligand-gated channels and second-scale photocycle (Kovalev et al., 2019). The broad representation of a distinct group of rhodopsins in virus genomes implies an important light-dependent function in virus-host interactions but the nature of this function remains obscure. Given that NCLDV play a major role in marine algae population dynamics, elucidation of the virus-host interaction mechanisms would make an important contribution to better understanding the impact of viruses on global ecology and climate (Gómez-Consarnau et al., 2019; Short, 2012).

Here we present the results of a comprehensive structure-function study of two homologous proteins from the VR1 group, OLPVR1 and VirChR1. We show that unlike previously reported data (Needham et al., 2019), VirChR1 demonstrate sodium/potassium-selective channelrhodopsin activity when expressed in human neuroblastoma cells. Upon light absorption, VirChR1 depolarizes cell membranes to a level sufficient to fire neurons. This finding, together with the fact that, in contrast to the previously characterized ChRs, VirChR1 is not permeable for calcium ions, suggests that viral rhodopsins could become invaluable optogenetic tools. Following functional characterization, we crystallized and solved multiple structures of OLPVR1 that revealed unique structure-function features of viral rhodopsins.

### Metagenomic search for viral rhodopsins genes and sequence analysis

To obtain a comprehensive set of rhodopsins in the vast metagenomic sequence database produced by the *Tara Ocean* project, we compared 36 rhodopsin sequences representative of the previously identified major groups to the sequences of all open reading frames from *Tara Ocean* contigs. This search retrieved 2584 Type 1 rhodopsins of which 80 belonged to viral group I and 172 belonged to viral group II as confirmed by phylogenetic analysis that also supported the monophyly of all viral rhodopsins (Extended Data Figure 5). Consistent with the monophyly of viral rhodopsins and the separation of groups 1 and 2, examination of sequence alignments detected several amino acid motifs that partially differed between the two groups. The amino acid triad implicated in proton exchange with the retinal Schiff base (residues 85, 89 and 96 in the reference bacteriorhodopsin (Luecke et al., 1999)) had the form DTS/DTT in the VR1 subfamily and DTT/DSV in the VR2 subfamily. Group 1 viral rhodopsins are characterized by several fully conserved residues, such as S11, Q15, E51, K192, N193, N197 and N205 (according to OLPVR1 numbering, Extended Data Figure 1) that are mainly located in proximity to the retinal Schiff base. In addition, the VR1 group has a signature topological feature, namely, an extended TM4 helix that consist of it transmembrane (TM4) and membrane-associated parts (ICL2) and has not been previously observed in characterized microbial rhodopsins (Figure 1b). Despite the overall low structural similarity with cryptophyte cation-conducting channelrhodopsins (Figure 1d), viral rhodopsins from group 1 retain the two highly conservative glutamates in TM2 (E44 and E51 in OLPVR1, and E83 and E90 in *Cr*ChR2) that have been shown to be critical for *Cr*ChR2 ion channelling (Kuhne et al., 2019; Wietek et al., 2014). Detailed analysis of the amino acid conservation in the VR1 subfamily (Extended Data Figure 3) shows that the majority of the conserved residues form the interior of the protein and can be predicted to contribute to ion-channeling of viral rhodopsins. Therefore, viral rhodopsins from group 1 with described conservativity are likely to share ion-channeling activity and functional features, so we name VR1 family members as viral channelrhodopsins or VirChR family.

**Figure 1.**
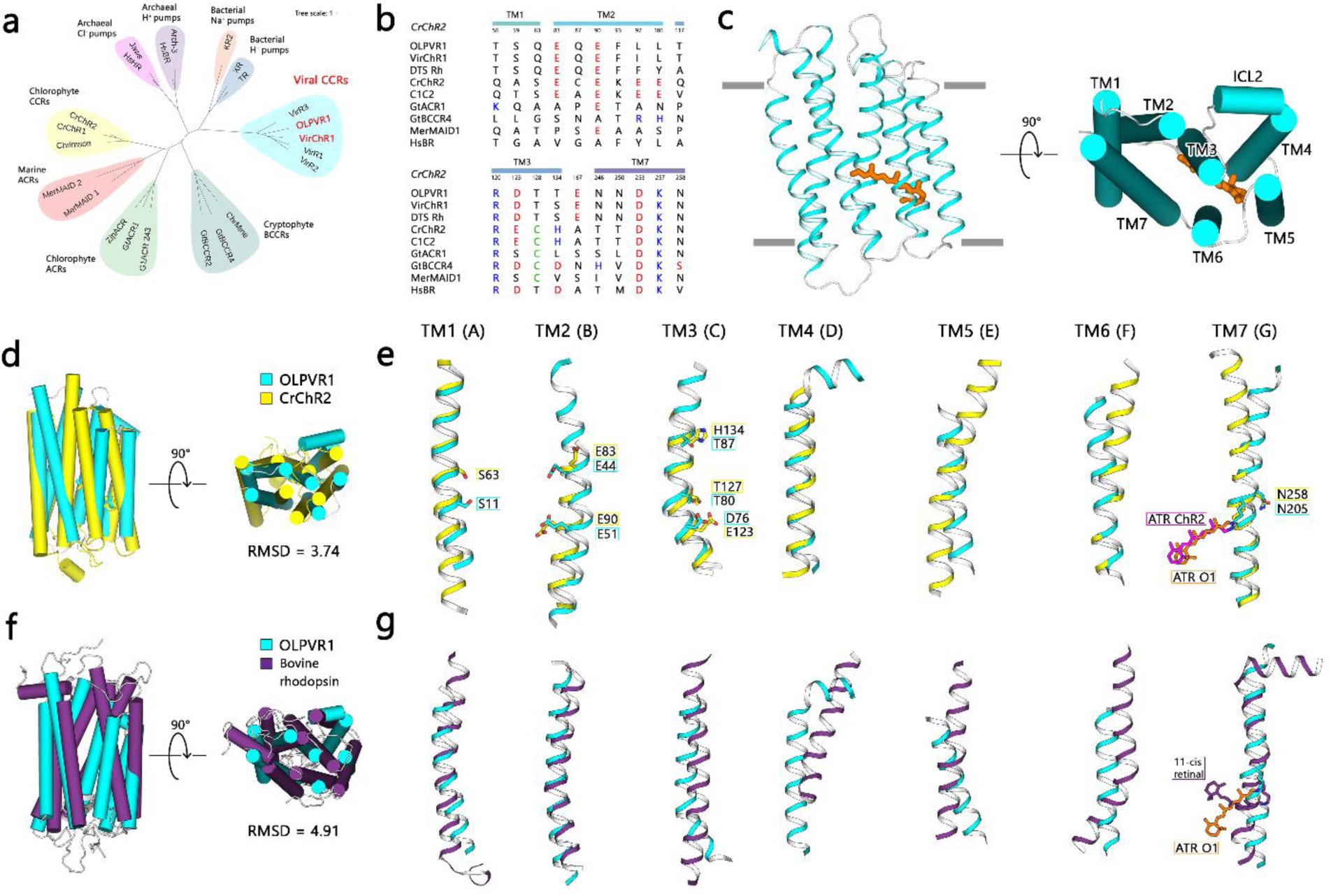
Phylogenetic and structural overview of the viral channelrhodopsins family. (a) Unrooted phylogenetic tree of the channelrhodopsin superfamily including viral channelrhodopsin representatives. Scale bar indicates the average number of amino acid substitution per site. CCR, cation-conducting channelrhodopsin, ACR, anion-conducting channelrhodopsin. OLPVR1 and VirChR1 proteins are additionally indicated in red. (b) Alignments of functionally important residues of transmembrane helices 1, 2, 3 and 7 of representative proteins from channelrhodopsin subfamilies. Positively and negatively charged residues are highlighted blue and red, cysteine residues are highlighted green (c) Crystal structure of OLPVR1 protein, viewed parallel to membrane (left) and from the extracellular side (right). All-trans retinal (ATR) is depicted with orange sticks. Rhodopsins were named according to the previous works (Govorunova et al., 2017; Klapoetke et al., 2014; Sineshchekov et al., 2017b). The hydrophobic membrane boundaries were calculated with the PPM server and are shown by gray lines (Lomize et al., 2012). (d) Structure alignment of OLPVR1 and *Cr*ChR2 (PDB: 6EID) structures colored cyan and yellow respectively. RMSD, root mean square deviation of atomic positions. (e) Individual TM helices are shown after superimposition of the OLPVR1 and *Cr*ChR2 rhodopsins. (f) Structure alignment of OLPVR1 and bovine rhodopsin (PDB: 1U19, (Okada et al., 2004)) structures colored cyan and purple respectively. (e) Individual TM helices are shown after superimposition of the OLPVR1 and bovine rhodopsin.

### Spectroscopic characterization of VirChR family

In order to characterize the molecular properties of viral channelrhodopsins, we expressed OLPVR1 and VirChR1 proteins in *E. coli*, and purified the proteins via a combination of affinity chromatography (Ni-NTA) and size-exclusion chromatography. Purified protein was incorporated into corresponding lipid-mimicking matrices and used for all spectroscopic experiments (see Methods for full details). UV-vis spectra of OLPVR1 and VirChR1 proteins reveal blue-light absorption with λ_max_ around 500 nm neutral pH values. Under acidic conditions the absorption spectra of both proteins undergo a reversible transition to the protonated state, associated with red-shifted absorption maximum at around 530 nm, similar to those for other microbial rhodopsins (Tsukamoto et al., 2013). In contrast to previously reported data, we did not observe any significant ion translocation ability of viral rhodopsins in pH change experiments with protein-containing liposomes (Needham et al., 2019). Under continuous bright light illumination, OLPVR1 liposomes have not showed any substantial pH change of the external solvent (Figure 2g). The maximum pH shift of OLPVR1 (0.03 pH units) is about 10 times less than that of the light-driven proton pump LR/Mac-containing liposomes. In order to elucidate photocycle kinetics of viral channelrhodopsins, we performed transient absorption measurements with OLPVR1-containing nanodiscs that revealed three distinct intermediate states of OLPVR1 photocycle. An early decaying K-like state (λ_max_ = 540 nm), followed by major accumulation of L-like (λ_max_ = 460 nm) and N-like (λ_max_ = 560 nm) states that live for about 1.5 seconds (Figures 2b and 2c). Unlike other channelrhodopsins, OLPVR1 lacks a detectable M-state, that is generally associated with the ion-conducting state of the protein (Kuhne et al., 2019). For viral channelrhodopsins, the equilibrium between L-like and N-like states is the major candidate for the ion-conducting state, where the structural changes in the protein may occur.

**Figure 2.**
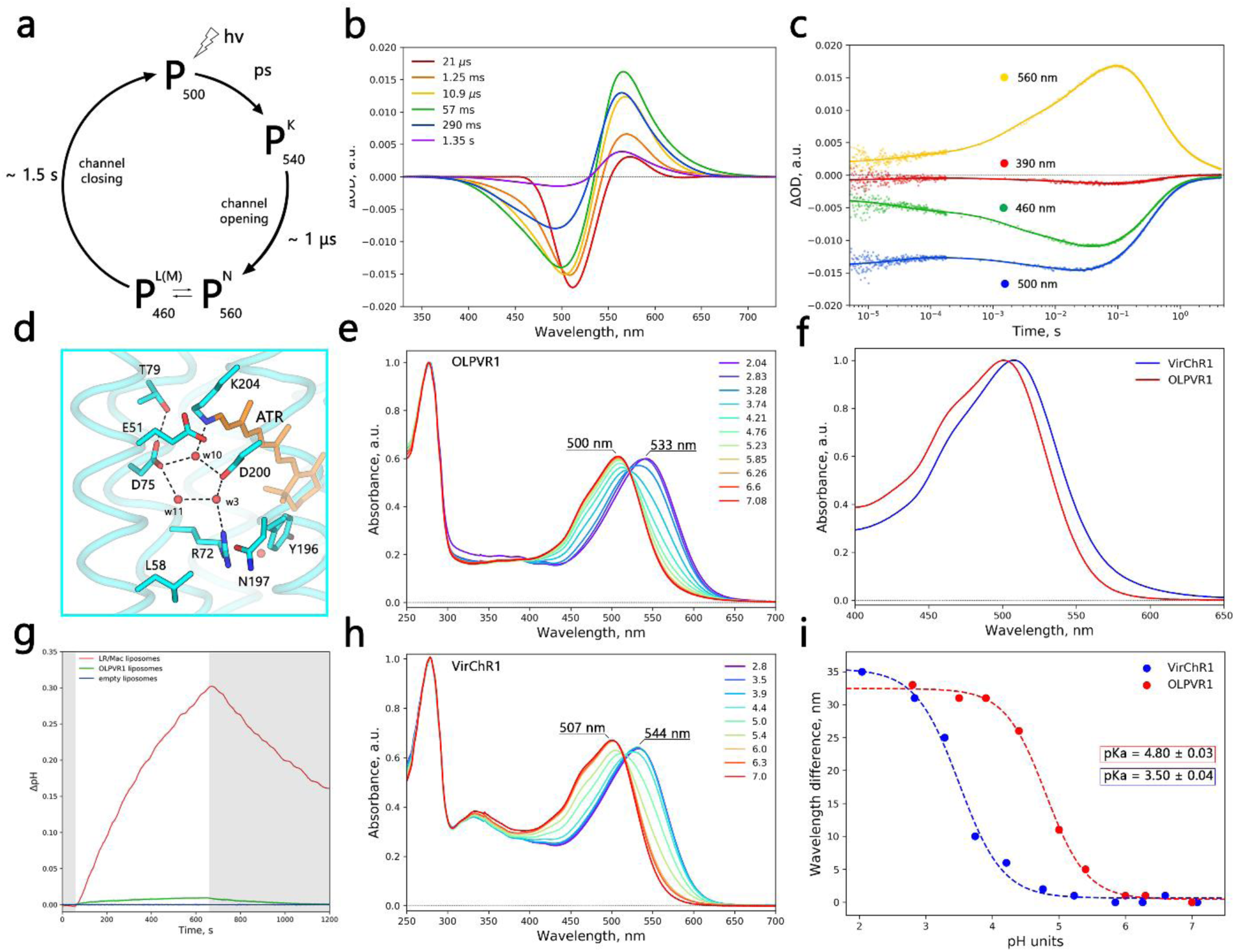
Spectroscopic characterization of viral channelrhodopsins. (a) Schematic model of viral rhodopsins photocycle. (b) Transient absorption spectra and (c) time traces at specific wavelengths of wild type OLPVR1 protein at pH 7.5. (d) Schiff base region of OLPVR1, important residues and water molecules are shown as sticks and spheres, hydrogen bonds are shown as dashed lines. (e) Absorption spectra of OLPVR1 at acidic (pH 2.8 - 7.0) pH range, normalized for absorption at 280 nm. (f) Normalized absorption spectra of OLPVR1 and VirChR1 proteins at neutral pH (pH 7.5). (g) Ion-transport activity assay of OLPVR1-containing proteoliposomes in 100 mM NaCl salt. Onset of illumination is indicated with white (light on) and gray (light off) background color, pH was adjusted to pH 6.0 prior to measurements. LR/Mac-containing liposomes and empty liposomes were used as positive and negative controls respectively. (h) Absorption spectra of VirChR1 at acidic (pH 2.8 - 7.0) pH range, normalized for absorption at 280 nm. (i) Red shift of UV-visible absorption spectrum and protonation of counterion of OLPVR1 and VirChR1. Wavelength maximum values are shown as circles. Sigmoidal curve fits are presented as dashed lines. The pKa values were calculated using sigmoidal fit.

### Electrophysiology of VirChR1, a light-gated cation channel

In order to identify the possible ion channeling activity, we expressed human codon-optimized OLPVR1 and VirChR1 genes, supplemented with RFP protein in SH-SY5Y human neuroblastoma cell line in the presence of all-trans retinal (see Methods for full details). Both proteins showed high expression levels in mammalian cells but OLPVR1 failed to localize to the plasma membrane according to the fluorescence microscopy and electrophysiology data. However, we observed robust expression of VirChR1 in neuroblastoma cells. Therefore, we characterized the ion channeling activity of VirChR1 as a representative of group1 viral rhodopsins. Consistent with high sequence similarity of OLPVR1 and VirChR1 (Figure 1c), we will later refer to the function of viral channelrhodopsins based on the data obtained for VirChR1 protein. Whole-cell patch clamp recordings reveal outwards currents generated by VirChR1 protein. Light-induced currents were observed in a bath solution of 10 mM HEPES pH 7.4, 110 mM NaCl, 2 mM MgCl_2_ and pipette solution of 10 mM HEPES pH 7.4, 110 mM NaCl, 2 mM MgCl_2_, 10 mM EGTA (hereafter both denoted standard In response to continuous illumination by LED light (λ_max_ = 470 nm) the photocurrents of VirChR1 revealed partial desensitization and photocurrents decay to a stationary level. Measuring photocurrents in different cells under standard conditions we did not see changes of kinetics or shifts of reversal potential. For the one representative neuroblastoma cell photocurrent stabilized at 50 pA at 80 mV (Figure 3a). The current values varied for different cells depending on the size of cell and protein expression level, but characteristic curve shape remained the same. During the measurement we observed the photocurrent level and its direction to depend on the membrane potential, meaning that light triggers passive ion conductance of VirChR1. VirChR1 exhibits an action spectrum similar to those of typical rhodopsins, with maximum sensitivity close to 500 nm (Figure 3d). Thus, both OLPVR1 and VirChR1 show retinal spectrum activation by blue light which is consistent with the fact that blue light penetrates more throughout the seawater photic zone(Jorgensen et al., 1987).

**Figure 3.**
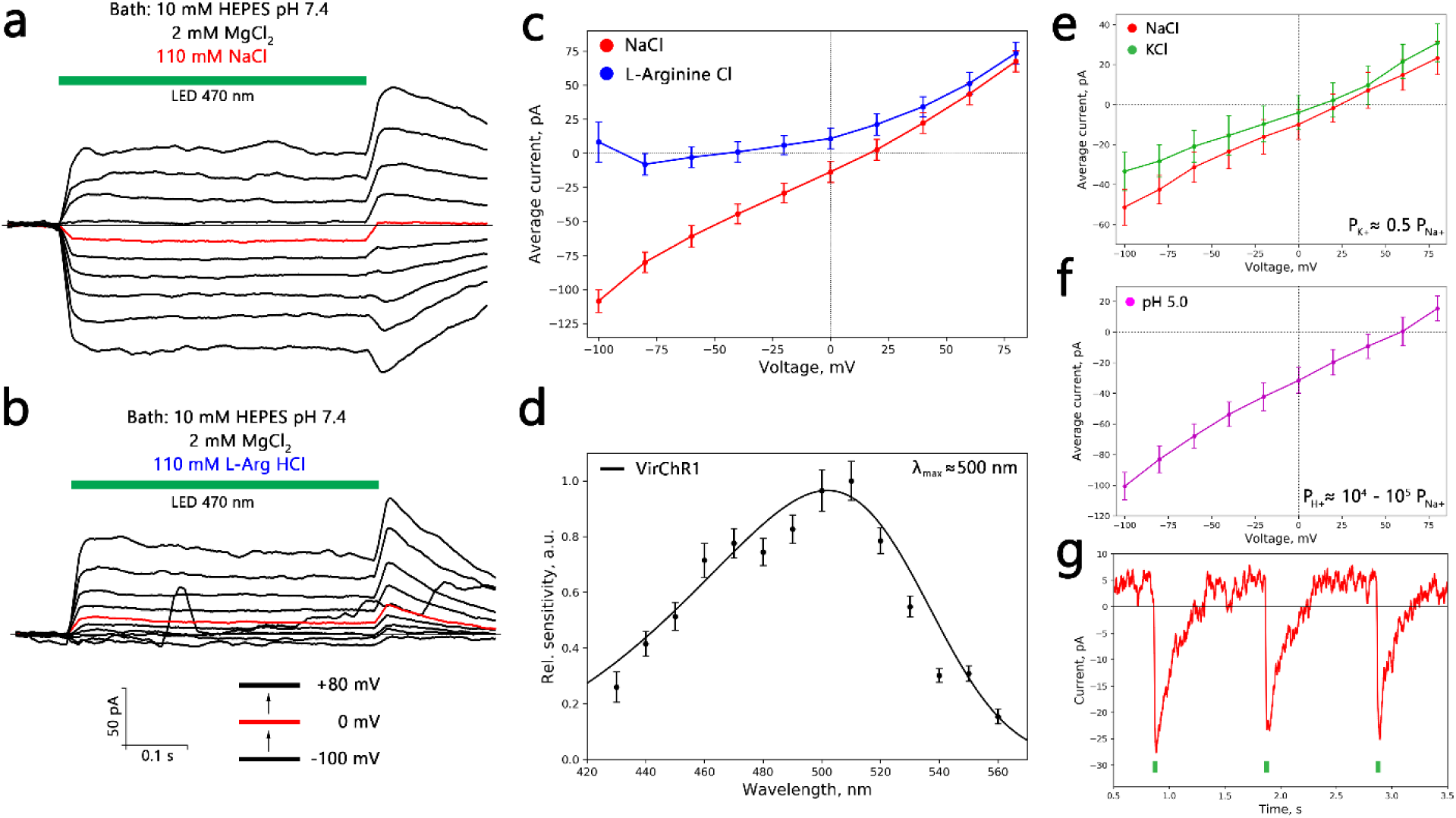
Ion selectivity and physiological features of VirChR1. Voltage-clamp records from one representative SH-SY5Y cell, expressing VirChR1 with (a) 10 mM HEPES pH 7.4, 110 mM NaCl, 2 mM MgCl_2_ and (b) 110 mM L-Arginine hydrochloride replacing NaCl in bath. Pipette solution during experiments was: 10 mM HEPES pH 7.4, 110 mM NaCl, 2 mM MgCl_2_, 10 mM EGTA. Currents are reproducible and typical to those in 30 experiments with other cells, illumination by LED (470 nm) lamp is indicated with green line. (c) Current-voltage dependences for one representative SH-SY5Y cell in 110 mM NaCl (red) and 110 mM L-Arginine hydrochloride (blue). (d) Action spectrum of VirChR1 measured using equal photon fluxes. (e-f) Current-voltage dependences for different bath/pipette solutions. Estimation of relative conductivities for different ions was done fitting traces with Goldman-equation. Traces are shown for (e) bath solutions: 110 mM NaCl (red) and 110 mM KCl (green) (pipette solution is standard) and for (f) pipette solution 110 mM L-Arginine hydrochloride salt solution of pH 5.0 (bath solution is standard). (g) Excitation recovery of photocurrent after short pulse of nanosecond laser (500 nm) activation.

Next, we performed ion substitution experiments in order to discriminate between possible cation and anion conductivity of the VirChR1 channel. First, we replaced standard bath solution with 63 mM Na_2_HPO4/NaH_2_PO4 pH 7.4 and 2 mM MgCl_2_ and found that this modification did not change neither the magnitude nor the reversal potential of the photocurrent. In contrast, when replacing 110 mM NaCl in bath solution with 110 mM L-Arginine hydrochloride, the inward current became immeasurably low, resulting in a dramatic change in the current-voltage dependence (Figure 3b). These results indicate that viral channelrhodopsins possess only cation-conducting activity.

### Unusual Ca^2+^ sensitivity of VirChR1

To evaluate the conductance of different cations by viral channelrhodopsins, we measured photocurrent dependencies on voltage in different bath solutions. Replacing Na^+^ with K^+^ cations (Figure 3e) in the bath solution yields an estimate of potassium permeability, P_K+_ ≈ 0.5·P_Na+._ To estimate H^+^ permeability, we replaced the standard bath solution with 10 mM Citric Acid pH 5.0, 110 mM L-Arginine hydrochloride, 2 mM MgCl_2_. Under these conditions, we observed full suppression of the photocurrent which occurred, presumably, due to the protonation of the Schiff base proton acceptor, similar to the case of OLPVR1 (Figure 2i). Thus, we replaced the pipette solution with 10 mM Citric Acid pH 5.0, 110mM L-Arginine hydrochloride, 2 mM MgCl_2_, 10 mM EGTA (Figure 3f), which restored the photocurrent to a measurable level. Fitting the photocurrent data with the Goldman-Hodgkin-Katz equation allows accurate estimation of H^+^ permeability, P_H+_ ≈ 10^4^ – 10^5^·P_Na+_. Overall, VirChR1 shows ion selectivity values comparable to those of *Cr*ChR2, namely, P_K+_ ≈ 0.5·P_Na+._ and P_H+_ ≈ 10^6^·P_Na+_. Thus, group 1 viral rhodopsins and cryptophyte cation channels are similar with respect to the conductivity of monovalent ions.

Next, we tested whether VirChR1 was permeable for divalent cations, such as Ca^2+^, similarly to *Cr*ChR2(Nagel et al., 2003). In fact, replacement of 110 mM NaCl in bath solution for 80 mM CaCl_2_ completely abolished the photocurrent (Figure 5a, 5b). The current-voltage dependences of VirChR1 in the presence and absence of Ca^2+^ indicate that VirChR1 is completely impermeable for Ca^2+^ cations (Figure 4c), in a sharp contrast to the high Ca^2+^ conductivity of *Cr*ChR2. Importantly, VirChR1 is fully blocked for both inward and outward ionic fluxes when concentration of Ca^2+^ exceeds a certain threshold. To characterize the kinetics of VirChR1 inhibition by Ca^2+^ ions, we measured dependence of the average photocurrent at +80 mV voltage on the CaCl_2_ concentration. A sigmoid-like dependence was identified, with the inflection point at K_Ca2+_ = 2.2 mM CaCl_2_, which is surprisingly close to the Ca^2+^ concentration in the world ocean (Tsunogai et al., 1973), and thus, suggestive of a functional role of viral rhodopsin inhibition by Ca^2+^ ions. Taken together, our findings suggest that VirChR1 is a light-gated cation channel that conducts exclusively monovalent ions (H^+^, Na^+^, K^+^) and is completely inhibited by divalent ions (Ca^2+^), with a characteristic enzyme-substrate kinetics (Figure 4f).

**Figure 4.**
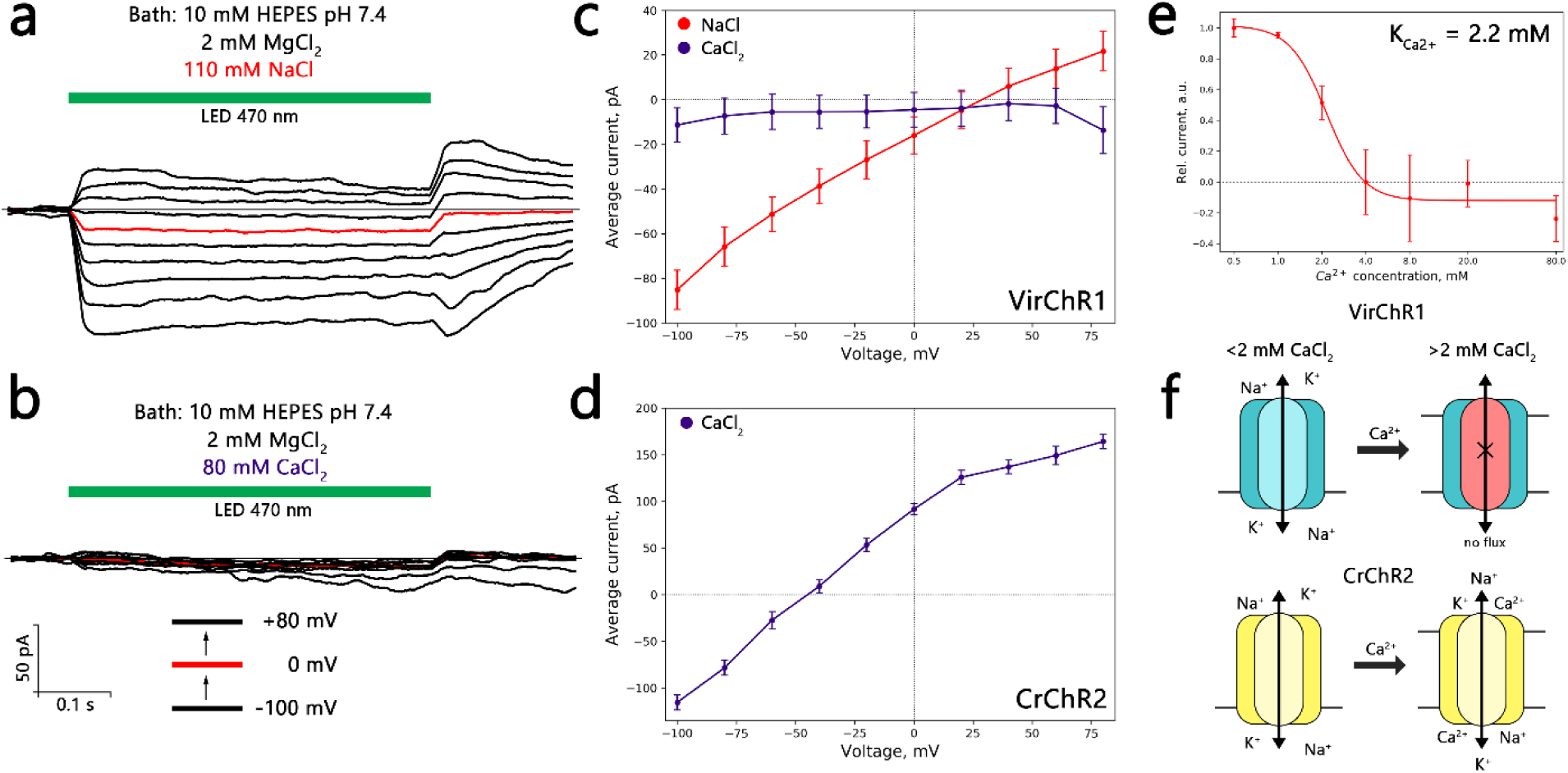
Calcium sensitivity of VirChR1 channel. Voltage-clamp records from one representative SH-SY5Y cell expressing VirChR1 in bath solution (a) 10 mM HEPES pH 7.4, 110 mM NaCl, 2 mM MgCl_2_ and (b) in 80 mM CaCl_2_ replacing NaCl in bath solution. Pipette solution: 10 mM HEPES pH 7.4, 110 mM NaCl, 2 mM MgCl2, 10 mM EGTA, illumination by LED (470 nm) lamp is indicated with green line. (c) Current-voltage dependences for one representative SH-SY5Y cell in 110 mM NaCl (red) and 80 mM CaCl_2_ (indigo) solutions. (d) Current-voltage dependences of a *Cr*ChR2-expressing SH-SY5Y cell in 80 mM CaCl_2_ solution. Current-traces had same shape in 3 more cells. Calcium blocking in CrChR2 was never observed in any cell. (e) Current dependence on calcium concentration in bath solution measured at +80 mV (inflection point is at approximately 2.2 mM of calcium). (f) Schematic comparison of VirChR1 and *Cr*ChR2 ion channeling activity under different calcium concentrations, proteins are colored cyan and yellow respectively.

**Figure 5.**
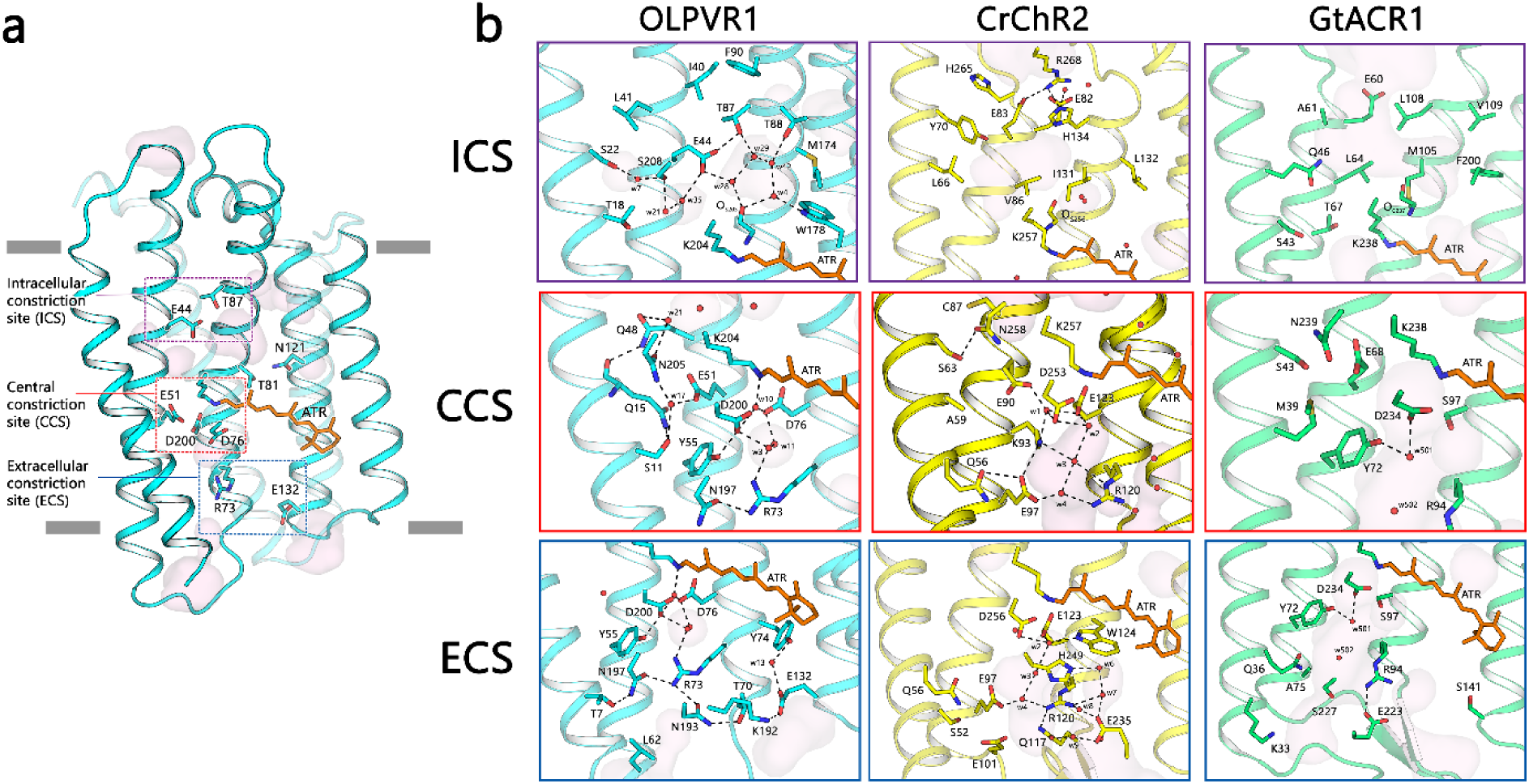
Organization of ion pathway constriction sites (CSs) in OLPVR1. (a) Three CSs and cavities forming the putative ion-conductive pathway in viral CCRs and highly conservative residues of OLPVR1. TM6 and TM7 helices are omitted for clarity. (b) Magnified view of the CSs in OLPVR1 (left), *Cr*ChR2 (middle, PDB: 6EID) and (right, PDB: 6CSN) structures, colored cyan, yellow and green respectively. Water accessible cavities were calculated using HOLLOW and are presented as pink surface (Ho and Gruswitz, 2008).

### Crystal structure of the viral rhodopsin

To decipher the molecular mechanism of ion channeling, the structure of viral channelrhodopsin from group 1 is essential. One crystal structure of the DTS-motif rhodopsin from *Phaeocystis globosa* virus 12T was recently reported(Needham et al., 2019). However, from the only one available structure it is not possible to distinguish the features of the entire group. Here, we present a high-resolution structure of another viral rhodopsin from group 1 OLPVR1 at 1.4 Å resolution. The protein was crystallized with an *in meso* approach similar to that used in our previous studies(Gushchin et al., 2015; Shevchenko et al., 2017). We obtained three different types of crystals. Type A rhombic crystals were grown at pH 8.0 using the monopalmitolein (MP) host lipid matrix and have the P2_1_2_1_2 space group with one protein molecule in the asymmetric unit. Type B hexagonal crystals were grown at pH 7.0 using a monoolein lipid matrix, have the P1 space group and contain two protein molecules in the asymmetric unit. OLPVR1 molecules are nearly identical in both structures (root mean square deviation (RMSD) less than 0.2 Å), so hereafter, we refer to the structure from type A crystals as it has the highest resolution. The crystal packing and examples of the electron density maps are shown in Extended Data Figures 7-8.

The structure of OLPVR1 protomer is composed of 7 transmembrane helices (TM1-7), connected by three intracellular and three extracellular loops. The OLPVR1 protein (residues 2-223), all-*trans* retinal (ATR) covalently bound to K204 (K257 in *Cr*ChR2 (Volkov et al., 2017)), 9 lipid molecules and 107 water molecules are clearly resolved in the electron density maps. Despite the fact that only one OLPVR1 protomer is present in the asymmetric unit, the crystal packing of the protein shows that OLPVR1 could be organized into homodimers, similar to those of *Cr*ChR2 (Kato et al., 2012; Volkov et al., 2017). These dimers might reflect the oligomeric state of the viral channelrhodopsin in the cell membrane (Extended Data Figure 8). The interfacial interaction in the putative dimer occurs mainly in the TM4 helix and involves several non-conservative residues, namely E108-E108’, Y111-Y111’, F122-F122’ interactions. Overall, OLPVR1 backbone is tolerantly superimposed with that of the Med12 proteorhodopsin (PDB ID: 4JQ6 (Ran et al., 2013)) with RMSD of 2.1 Å, whereas the alignments with the *Hs*BR (PDB ID: 1C3W (Luecke et al., 1999)) and *Cr*ChR2 (PDB ID: 6EID) structures gives RMSD values of 4.3 and 3.74 Å, respectively (Figure 1d, Extended Data Figure 6). Interestingly, unlike other microbial rhodopsins, OLPVR1 architecture closely resembles the architecture of G-protein coupled receptors with TM3 helix protruding to the center of the protein. In particular, OLPVR1 align with bovine rhodopsin (Okada et al., 2004) with RMSD of 4.91 Å (Figure 1f), with a high similarity among helices forming ion conducting pathway (TM1-3 and TM7).

OLPVR1 protomer has short extracellular loops which sharply differentiates it from other channelrhodopsins, that typically have large N- and C-terminal domains. Unlike in other microbial rhodopsins, helices TM3 and TM4 of OLPVR1 are connected by the loop containing the membrane-associated helix (ICL2 helix) which is composed of hydrophilic residues. Strikingly different from other rhodopsins, the intracellular parts of TM6 and TM7 helices of OLPVR1 are significantly moved apart from each other far enough to form a pore (Extended Data Figure 6). The pore is located at about 8 Å from the cytoplasmic side of the lipid membrane border (Extended Data Figure 3a) and connects the inside of the protein with the groove on its surface, which leads further to the intracellular bulk. Surprisingly, in our structure, the groove and a part of the pore are occupied with the fragment of MP molecule, a host lipid of the crystallization matrix, which is likely to be a crystallization artifact and is discussed in more details in the Supplementary Text.

### Structure of the retinal binding pocket and RSB region

The retinal cofactor is covalently attached to the conserved K204 residue in OLPVR1. 2F_o_ – F_c_ electron density maps at 1.4 Å reveal two alternative conformations of the retinal in the region of β-ionone ring, while the configuration near the Schiff base in both of them is all-*trans*, 15-*anti*. The RSB region of OLPVR1 retinal binding pocket is very similar to that of *Hs*BR (Extended Data Figure 9c). Particularly, the D75 and D200 side chains and water molecules w3, w10 and w11 in OLPVR1 (corresponding to D85, D212, w401, w402 and w406 in *Hs*BR, respectively) form an almost identical to *Hs*BR pentagon-like hydrogen bonds structure, which is important for proton transfer in light-driven proton pumps (Freier et al., 2011). Moreover, the pentagon is similarly stabilized by T79 and R72 (T89 and R82 in *Hs*BR, respectively). In contrast, the pair of RSB counterions in CrChR2 is composed of E123 and D253 which, together with water molecules positioned in this region, presumably results in notably different stabilization of the Schiff base as compared to OLPVR1 (Extended Data Figure 10). The walls of the retinal pocket of OLPVR1 around the polyene chain are composed of several aromatic amino acids similar to those in *Hs*BR, namely W87, W178 and Y181 (Extended Data Figure 10a). However, there are several important changes near the β-ionone ring, particularly, L142, G182 and F185 instead of W138, P186 and W189 in *Hs*BR. These amino acids can potentially be candidates for mutation scanning in order to obtain red-shifted versions of VirChRs, more suitable for practical applications, considering that OLPVR1 and *Hs*BR have retinal absorption maxima at 500 nm and 560 nm respectively.

A characteristic feature of the retinal binding pocket of OLPVR1 (and, presumably, all viral channelrhodopsins) is the presence of highly conserved and directly hydrogen bonded residues T81 and N121 at the positions of the corresponding residues T80 and D115 in *Hs*BR, and C128 and D156 in *Cr*ChR2 (Extended Data Figure 10b). These pairs connect the middle parts of TM3 and TM4. Importantly, in case of ChChR2, where C128 and D156 are interconnected by hydrogen bonds via a water molecule, the alteration of the pair dramatically affects the kinetics of the protein and it was suggested that the pair is involved in the RSB reprotonation during photocycle(Berndt et al., 2009; Nack et al., 2010; Volkov et al., 2017)

### Organization of the OLPVR1 ion-conducting pathway

The structure suggests that the ion-conducting pathway of OLPVR1 is formed by TM1-3 and TM7 helices and is lined with several water-accessible cavities (Figure 5a). Unlike other channelrhodopsins, OLPVR1 lacks any prominent cavities in the extracellular part of the protein, but instead has a pore in the intracellular part, which ends up with a relatively large hydrophilic cavity inside the protein near the retinal (Figure 4a, b). The cavity is filled with four water molecules (w4, w28, w29, w42) and surrounded by polar residues E42, T87, T88 and W178. Water molecules together with the backbone oxygen of S203 residue form a hydrogen bond pentagon (Figure 5b) and may play a role in the hydration of cation during its translocation. A dense hydrogen bonding network involving water molecules and polar/charged residues protrude from the cavity almost to the extracellular bulk, only breaking in the central region near E51 residue. The putative ion pathway includes three constriction sites inside the protein (Extended Data Figure 12). Each site (described in details below) is comprised of highly conserved residues (Extended Data Figure 3). The constriction sites regions are almost identical in OLPVR1 and DTS-motif rhodopsin from *Phaeocystis globosa* virus 12T (Needham et al., 2019), therefore we consider them as characteristic feature of the group 1 of viral rhodopsins. Hence, we assume that they are also essential for the ion channeling.

The intracellular constriction site (ICS) of the OLPVR1 is relatively loose and formed mostly by E44 side chain. It separates the large intracellular cavity from a polar region near T18, S22, Q48, N205 and S208 in the middle part of the protein, containing also three water molecules w7, w21 and w35. E44 side chain is pointed towards the retinal, similarly to E122 in C1C2 (Kato et al., 2012) (Extended Data Figure 12), and is stabilized by hydrogen bonds with T87 and water molecules w20 and w35. Interestingly, unlike in CrChR2 and Chrimson (Oda et al., 2018), where the intracellular constriction sites (intracellular gates) are almost 14 Å far from the RSB and separated from it by the hydrophobic cavity, the ICS of OLPVR1 is located closer (9 Å) to the active center and is connected by extended hydrogen bonding network to the central constriction site (CCS). Moreover, in CrChR2 and Chrimson the ICS is comprised of tightly connected charged amino acids, completely blocking the flow of ions in the resting state. In these terms, the lack of compaction in the cytoplasmic region of OLPVR1 and the existence of the pore between TM6 and TM7 make the organization of the intracellular part of the protein closer to that of anion channel GtACR1(Kim et al., 2018), where the pore protrudes from the intracellular bulk almost down to the retinal without any constrictions (Figure 5d). This might mean a different gating mechanism in the cytoplasmic part of OLPVR1 from most commonly used light-driven cation channels.

The CCS of the OLPVR1 includes, fully conserved among VirChR group, S11, Q15, E51 and N205 residues that likely hinder ion translocation pathway in the resting state (Figure 6a). The core of the CCS is comprised of S11-E51-N205 (S-E-N) triad, which is similar to S63-E90-N258, S102-E129-N297 and S43-E68-N239 clusters of central gates (CGs) in CrChR2, C1C2 and GrACR1, respectively (Figure 6d). However, unlike in other channelrhodopsins, in OLPVR1 these residues are interconnected by the water molecule (w17), which thus define their orientation, even though no water accessible cavities were predicted in this region (Figure 5b). Moreover, S11 is located one α-helix turn closer to the extracellular side than the corresponding serines in CrChR2, C1C2 and GtACR1 and is additionally stabilized by the hydrogen bond with the Q15 side chain. Q15 is a unique residue in OLPVR1 and has no analogues in other channelrhodopsins. In general, the organization of the S-E-N triad in OLPVR1 is closer to that of C1C2 and GtACR1, but not CrChR2 (Figure 4), where E90 side chain is pointed towards the RSB region and is a part of the extended hydrogen bond network in the extracellular part of the protein. Thus, unlike in CrChR2, in OLPVR1 the CCS (and particularly E51) is not directly connected to the RSB region in the resting state.

**Figure 6.**
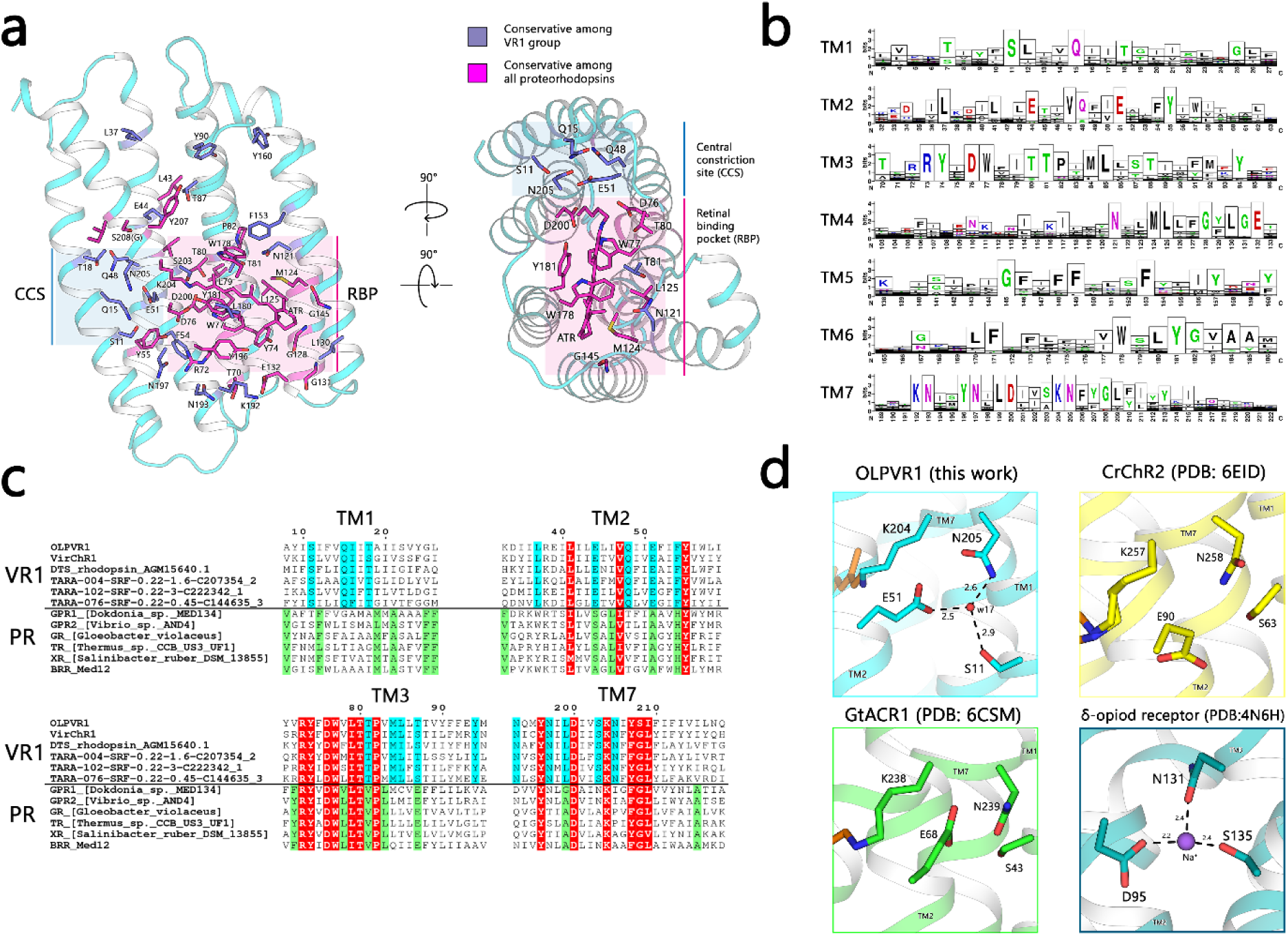
Conservativity analysis of viral rhodopsins. (a) Structural overview of highly conservative (70% cutoff) residues among viral rhodopsins family (n = 124). OLPVR1 structure was used as a template of viral rhodopsin, viewed parallel from membrane (left) and from the cytoplasm (right). Residues conservative among all proteorhodopsins and residues exclusively conservative by viral rhodopsins are shown as sticks and colored magenta and dark blue respectively. (b) Sequence logo of transmembrane helices (TM1 – TM7) of viral rhodopsins family created using Weblogo sequence generator server(Crooks, 2004). (c) Sequence alignment of TM1, TM2, TM3 and TM7 helices of 6 representative sequences from viral rhodopsins and marine proteorhodopsins. Residues conservative among VR1, PR, and both groups are colored cyan, greed and red respectively. (d). The magnified view of the highly conservative S-E-N triad that comprise central constriction site (CCS) in OLPVR1 (top left), CrChR2 (PDB: 6EID, top right), and GtACR1 (PDB: 6CSM, down left). The sodium binding site formed by S-D-N triad in human delta-opioid receptor (PDB: 4N6H (Fenalti et al., 2014), down right).

The extracellular constriction site (ECS) of OLPVR1 includes highly conserved R72, E132, K192, N193 and N197 residues that are tightly interconnected by hydrogen bonds. In contrast to extracellular gates (ECGs) of CrChR2 and GtACR1, R72 side chain is oriented towards the RSB, similarly to the analogous R82 in HsBR, however is stabilized by highly conserved N193 and N197 residues, while R82 in HsBR is stabilized mostly by the surrounding water molecules(Freier et al., 2011; Luecke et al., 1999). Also, unlike CrChR2, OLPVR1 lacks polar/charged amino acids in the region of E97 and E101 at the TM2 of CrChR2, which are substituted by leucines in the viral channelrhodopsin. Besides that, OLPVR1 possesses a more compact configuration of residues in the extracellular part than CrChR2, resulting in a more confined water-accessible cavities architecture of OLPVR1. This might be another reason why virus channelrhodopsins from group 1 are not permeable for larger ions.

## Discussion

### Distinct structural features of group 1 viral rhodopsins

Integration of electrophysiological, spectroscopic and structural analyses reveal unique features of group 1 viral rhodopsins as compact, Ca^2+^-inhibited channelrhodopsins. Given the diversity of viral rhodopsins, we can infer features only for the subset of group 1 channelrhodopsins that share a set of highly conserved residues, and specifically S-E-N triad (S11, E51 and N205) that are likely to play a key role in the ion-conducting mechanisms of viral channelrhodopsins. To the best of our knowledge, the exclusive conductivity for monovalent cations and its regulation by divalent cations have not been reported for any rhodopsins. Viral channelrhodopsins have proteorhodopsin-like architecture with short extracellular loops and share several structural features, such as membrane-associated ICL2 helix and an unconventional TM6-TM7 orientation. In a stark contrast to *Cr*ChR2 and other CCRs, viral channelrhodopsins share a set of highly conserved residues that encompass the ion-conducting pathway and, by analogy with *Cr*ChR2, possess three consecutive gates that are likely to displace in the open state of the channel. Notably, viral channelrhodopsins lack the DC amino acid pair (C128 and D156 in *Cr*ChR2), which is replaced by a NT pair (T88 and N121 pair in OLPVR1) that is conserved in all group 1 viral rhodopsins. Overall, this study provides a wide general insight into functional and structural features of viral channelrhodopsins although additional research is clearly needed to fully characterize their biochemical activity and biological function.

### Predicted biological role of viral channelrhodopsins

In this work, we demonstrate that group 1 viral rhodopsins are Na^+^/K^+^ selective light-gated ion channels that are inhibited by divalent cations. The fact that viral channelrhodopsins are widely distributed across the world ocean, where they are primarily constrained to the photic zone, offers clues to their likely biological role. The NCLDV-encoded group 1 viral rhodopsins infect microalgae, a key component of the phytoplankton (Uitz et al., 2010; Yau et al., 2011). What could then be the function of virus-encoded ion channels when expressed in the infected host? A plausible answer to this question is suggested by the analogy with the well-studied *Cr*ChR1 and *Cr*ChR2 channelrhodopsins from chlorophyte alga *Chlamydomonas reinhardtii* (Sineshchekov et al., 2002). These channelrhodopsins have been shown to sense ambient light and to transmit the signal downstream to the flagellar motor of motile algae, allowing them to migrate to an environment with light conditions optimal for photosynthesis and survival (Harz and Hegemann, 1991). The primary function of *Cr*ChR1 and *Cr*ChR2 is activation of secondary Ca^2+^-channels by changing the electrical polarization of the cell which mediates phototaxis and photophobic responses (Nultsch et al., 1986; Sineshchekov et al., 2009). Photoexcitation of the channelrhodopsin receptors results in generation of photoreceptor currents and membrane depolarization followed by activation of voltage-gated calcium channels triggering the flagellar motion (Böhm et al., 2019; Jékely, 2009; Kianianmomeni et al., 2009; Sineshchekov et al., 2002). Recently, the cryptophyte alga *Guillardia Theta* has been reported to possess genes for three distinct groups of channelrhodopsins, namely, cation- and anion- conducting channelrhodopsins with high similarity to chlorophyte channelrhodopsins, and a third group of bacteriorhodopsin-like channelrhodopsins (BCCRs) with an unconventional architecture (Govorunova et al., 2016; Sineshchekov et al., 2017a). Surprisingly, BCCRs are inhibited by high concentrations of calcium (greater than 40 mM CaCl2), suggesting a functional similarity with viral rhodopsins (Shigemura et al., 2019). Given that large and especially giant viruses possess diverse genes that enhance the host metabolic functions and hence promote virus reproduction (Lindell et al., 2005; Sharon et al., 2009; Filée, 2018), it seems likely that viral rhodopsins function as additional ion channels to supplement and augment the host phototactic systems similarly to the role of BCCRs in cryptophyte algae and thus boost metabolic processes required for virus reproduction. Although it remains unclear whether BCCRs are also inhibited by physiological concentrations of calcium, it appears likely that, due to their impermeability to Ca^2+^, viral channelrhodopsins and, presumably, also BCCRs activate secondary Ca^2+^-channels only by membrane depolarization, but not via biochemical amplification, as has been suggested for *Cr*ChR1 and *Cr*ChR2 (Sineshchekov et al., 2009).

Phytoplankton is the base of the marine food chain and an important source of carbon in the global carbon cycle(Charlson et al., 1987). Therefore, the algal population makes a major contribution to the regulation of many aspects of the global environment, and in particular, climate control (Moore et al., 2008). Viral rhodopsins are likely to play an important role in the regulation of phytoplankton dynamics by NCDLV.

### Electrophysiological features of VirChR1 and potential utility of viral channelrhodopsins for optogenetic applications

The results of this work (for example, Fig. 3a) demonstrate photocurrent overshooting after turning the light down. Although the cause of this remains unknown, we suggest that it indicates second-photon absorption during continuous light measurements. Also, non-zero negative photocurrent was observed under symmetrical conditions at 0 mV (Fig 3c, it also results in positive reversal potential). The same negative photocurrent was also detected when channel activity was blocked by calcium (Fig 4c, 4e). We suggest that inward-pumping activity of VirChRs is responsible for this current.

The finding that, in contrast to *Cr*ChR2 and other known CCRs, viral channelrhodopsins are impermeable to divalent cations, such as Ca^2+^, suggest that VirChRs might contribute to a new generation of optogenetic tools (Renault et al., 2015; Stamatakis et al., 2018). In particular, because their application would not interfere with important native Ca^2+^-dependent processes in the cells. Ca^2+^-impermeable channelrhodopsins could become invaluable tools to study processes in the brain, where optogenetic regulation of synapses is required (Lin, 2011; Lin et al., 2013; Tritsch et al., 2016).

## Methods

No experiments in animals were conducted in this paper and hence experiments were not randomized or blinded.

### Metagenomic analysis

Group I viral rhodopsins, were retrieved from metagenomic assembled contigs through combining similarity search, protein clustering and Hidden Markov Models. Briefly, a first dataset of *bona fide* rhodopsins was retrieved from *TARA Ocean* metagenome assembled contigs that were downloaded from ENA (https://www.ebi.ac.uk). Coding DNA sequences were predicted from contigs longer than 2 Kb using Prodigal (Hyatt et al., 2010), and annotated against the NR database of NCBI using Diamond (Buchfink et al., 2015). *Bona fide* rhodopsins were selected by screening for different keywords related to rhodopsins that must be contained in the annotation, and furthermore filtered according to 7 transmembrane domains that were predicted using Phobius(Käll et al., 2004). Selected proteins were next aligned using the R package Decipher (Wright, 2015), and alignments were used to infer a phylogenetic tree through FastTree (Price et al., 2010) and using default parameters. Phylogenetic distances between nodes on the tree topology were considered in order to cluster *bona fide* rhodopsins into distinct clades, each of which was used to train a Hidden Markov Model (HMM). All HMMs were finally queried against *TARA Ocean* assembled contigs using HMMER version v3.1b2 (http://hmmer.org) and setting an e-value threshold of 1e-5. Proteins identified through HMMs were clustered at 100% identity using CD-HIT suite(Li and Godzik, 2006) in order to remove redundancy, and reduced to a total of 2584 Type-1 rhodopsins that were further analysed.

The dataset of group I viral rhodopsins was constructed by searching the NCBI non-redundant protein sequence databases along with the TARA metagenomic sequences using BLSTP and TBLASTN. For the sake of clarity, for Extended Data Figure 1 we used a reduced number of sequences. To obtain a representative set of 16 TARA metagenomic sequences with OLPVR1 and VirChR1 sequences included, we used the CD-HIT suite□ with default parameters and 60% identity cut-off.

### Sequence alignment and phylogenetic analysis

Rhodopsin sequences were aligned using MUSCLE using UGENE software (Okonechnikov et al., 2012) with the default parameters. Type-1 rhodopsins were named according to their names in literature. The sequence alignment was created using ESPript3 online server (Robert and Gouet, 2014). Phylogenetic tree reconstruction was conducted by PHYLIP Neighbor Joining method using UGENE software (Okonechnikov et al., 2012) with the following parameters: Jones-Taylor-Thornton model, transition/transversion ratio = 2.0, no gamma distribution applied. Tree visualization was done using iTOL server (Letunic and Bork, 2016). GenBank accession numbers are additionally indicated.

### Cloning, expression, and purification

OLPVR1-coding DNA was synthesized commercially (Eurofins). The nucleotide sequence was optimized for E. coli expression using the GeneOptimizer software (Life Technologies). The gene, together with the 5′ ribosome-binding sites and the 3′ extensions coding additional LEHHHHHH* tag, was introduced into the pEKT expression vector (Novagen) via NdeI and XhoI restriction sites. The plasmid was verified by sequencing. The protein was expressed as described previously (Shevchenko et al., 2017) with modifications. E. coli cells of strain C41 (StabyCodon T7, Eurogentec, Belgium) were transformed with the expression plasmid. Transformed cells were grown in shaking baffled flasks in an autoinducing medium ZYP-5052 containing 50 mg/L kanamycin at 37°C. When the OD_600_ in the growing bacterial culture is in 0.8 – 1.0 range (glucose level < 10 mg/L), 10 μM all-trans-retinal (Sigma-Aldrich) and 1 mM isopropyl β-d-1-thiogalactopyranoside were added, the incubation temperature was reduced to 20°C and incubated for 18 hours. After incubation cells were collected by centrifugation (4,500 rpm, 30 min) and disrupted in an M-110P Lab Homogenizer (Microfluidics) at 25,000 p.s.i. in a buffer containing 20 mM Tris-HCl, pH 8.0 with 50 mg/L DNase I (Sigma-Aldrich). The membrane fraction of the cell lysate was isolated by ultracentrifugation at 35000 rpm for 1 h at 4°C (Type 70 Ti Fixed-Angle Titanium Rotor, Beckmann). The pellet was resuspended in a buffer containing 20 mM NaH_2_PO_4_/Na_2_HPO_4_, pH 8.0, 0.1 M NaCl and 1% n-Dodecyl β-D-maltoside (DDM, Anatrace, Affymetrix) and stirred for 18 hours for solubilization. The insoluble fraction was removed by ultracentrifugation at 35,000 rpm for 1 hour at 4°C. The supernatant was loaded on an Ni-NTA column (Qiagen), and washed with a buffer containing 10 mM NaH_2_PO_4_/Na_2_HPO_4_, 0.1 M NaCl, 0.01 M imidazole and 0.05% DDM buffer (pH 8.0). Elution of the protein was done in a buffer containing 10 mM NaH_2_PO_4_/Na_2_HPO_4_, 0.1 M NaCl, 0.5 M imidazole and 0.05% DDM (pH 8.0). The eluate was subjected to size-exclusion chromatography on a 20 ml Superdex 200i 10/300 GL column (GE Healthcare Life Sciences) in a buffer containing 10 mM NaH_2_PO_4_/Na_2_HPO_4_, pH 8.0, 0.1 M NaCl and 0.05% DDM. Protein-containing fractions with an A_280_/A_500_ absorbance ratio (peak ratio, p.r.) of lower than 1.5 were pooled and dialyzed against 100 volumes of 10 mM NaH_2_PO_4_/Na_2_HPO_4_, 0.1 M NaCl, and 0.05% DDM (pH 8.0) buffer twice for 2 hours to dispose imidazole. The purified protein was concentrated for 40 mg/ml for crystallization.

### Reconstitution of the protein into lipid-based systems

Phospholipids (azolectin from soybean, Sigma-Aldrich) were dissolved in CHCl_3_ (Chloroform ultrapure, Applichem Panreac) and dried under a stream of N_2_ in a glass vial. The solvent was removed by overnight incubation under vacuum. The dried lipids were resuspended in 100 mM NaCl buffer supplemented with 2% (w/v) sodium cholate. The mixture was clarified by sonication at 4°C and OLPVR1 was added at a protein/lipid ratio of 1:20 (w/w). The detergent was removed by 2 days stirring with detergent-absorbing beads (Amberlite XAD 2, Supelco). The mixture was dialyzed against 100 mM NaCl, (pH 7.0) buffer at 4°C for 8 hours to adjust the desired pH. The obtained liposomes were used for the measurement of pump activity with pH electrode. The OLPVR1-containing nanodiscs were assembled using a standard protocol described elsewhere (Ritchie et al., 2009)□. 1,2-dimyristoyl-sn-glycero-3-phosphocholine (DMPC, Avanti Polar Lipids, USA) and an MSP1D1 version of apolipoprotein-1 were used as a lipid and scaffold protein, respectively. The molar ratio during assembly was DMPC:MSP1D1:OLPVR1 = 100:2:3. The protein-containing nanodiscs were dialyzed against 100 volumes of 10 mM NaH_2_PO_4_/Na_2_HPO_4_, 100 mM NaCl (pH 7.5) buffer twice and then subjected to size-exclusion chromatography on a 20 ml Superdex 200i 10/300 GL column (GE Healthcare Life Sciences) for detergent removal.

### Ion-trafficking assay with protein-containing liposomes

The measurements were performed on 2 ml of stirred proteoliposome suspension at 0°C. OLPVR1- and LR/Mac- containing liposomes were prepared following the protocol described above. Liposomes were illuminated for 10 minutes with a halogen lamp (Intralux 5000-1, VOLPI) and then were kept in the dark for another 10 minutes. Changes in pH were monitored with a pH-meter (LAB 850, Schott Instruments). Some of the measurements were repeated in the presence of 30 μM of carbonyl cyanide m-chlorophenyl hydrazine (CCCP, Sigma-Aldrich) under the same conditions.

### pH titration

To investigate the pH dependence of the absorption spectra of OLPVR1, about 6 μM protein was suspended in the titration buffer (10 mM citrate, 10 mM MES, 10 mM HEPES, 10 mM MOPS, 10 mM CHES and 10 mM CAPS). Then, the pH was changed by addition of diluted or concentrated HCl or NaOH to obtain 0.5-0.7 pH change. The absorption spectra were measured with a UV-visible spectrometer (V-2600PC, Shimadzu).

### VirChR1 expression in SH-SY5Y cells

The human codon optimized VirChR1 gene was synthesized commercially (Eurofins). The gene was cloned into the pcDNA3.1(-) vector bearing an additional membrane trafficking signal, a P2A self-cleaving peptide, and a GFP variant from the C-terminal part of the gene, and Haemogluttinin and Flag-tag peptides from the N-terminal part of the gene(Gradinaru et al., 2010; Shcherbo et al., 2007). The SH-SY5Y human neuroblastoma cells at a confluency of 80-90% were transfected with the plasmid and Lipofectamine LTX according to the manufacturer’s protocol (Thermo Fisher Scientific). The cells were incubated under 5% CO_2_ at 37°C. After transfection (16-24 h), electrophysical experiments were performed.

### Electrophysiological recordings

For the electrophysiological characterization of VirChR1, whole-cell patch clamp recordings were performed (Scientifica LASU, Axon Digidata 1550A, Multiclamp 700B). Horizontal puller (Sutter Instrument CO, Model P-2000) was used for fabrication of patch pipettes (borosilicate glass GB150F-8P, 3 – 6 MΩ). Experiments were conducted using SH-SY5Y cell line. Photocurrents were measured in response to LED light pulses with saturating intensity λ = 470 ± 20 nm. For the action spectra ultrashort nanosecond light pulses were generated by Brilliant Quantel using OPO Opotek MagicPrism for different wavelenghts.

#### Time resolved absorption spectroscopy

Excitation/detection systems were composed as such: Brilliant B laser with OPO Rainbow (Quantel Inc.) was used, providing pulses of 4-ns duration at 530-nm wavelength and an energy of 2 mJ per pulse. Samples (5 × 5–mm spectroscopic quartz cuvette; Hellma GmbH & Co.) were placed in a thermostated house between two collimated and mechanically coupled monochromators (LOT MSH150). The probing light (xenon arc lamp, 75 W, Hamamatsu) passed the first monochromator sample and arrived after a second monochromator at a photomultiplier tube (PMT) detector (R12829, Hamamatsu). The current-to-voltage converter of the PMT determines the time resolution of the measurement system of ca. 50 ns (measured as an apparent pulse width of the 5-ns laser pulse). Two digital oscilloscopes (Keysight DSO-x4022A) were used to record the traces of transient transmission changes in two overlapping time windows. The maximal digitizing rate was 10 ns per data point. Transient absorption changes were recorded from 10 ns after the laser pulses until full completion of the phototransformation. At each wavelength, 25 laser pulses were averaged to improve the signal-to-noise ratio. The quasilogarithmic data compression reduced the initial number of data points per trace (∼32,000) to ∼850 points evenly distributed in a log time scale giving ∼100 points per time decade. The wavelengths were varied from 330 to 700 nm in steps of 10 nm using a computer-controlled step motor. Absorption spectra of the samples were measured before and after each experiment on a standard spectrophotometer (Avantes Avaspec 2048L). Obtained datasets were independently analyzed using the multiexponential least-squares fitting by MEXFIT software (Chizhov et al., 1996). The number of exponential components was incremented until the SD of weighted residuals did not further improve. After establishing the apparent rate constants and their assignment to the internal irreversible transitions of a single chain of relaxation processes, the amplitude spectra of exponents were transformed to the difference spectra of the corresponding intermediates in respect to the spectrum of the final state.

#### Crystallization

The crystals of OLPVR1 and O1O2 proteins were grown with an in meso approach(Caffrey and Cherezov, 2009), similar to that used in our previous works (Gushchin et al., 2015; Kovalev et al., 2019; Shevchenko et al., 2017). In particular, the solubilized protein (40 mg/ml) in the crystallization buffer was mixed with premelted at 50°C monoolein (MO, Nu-Chek Prep) or monopalmitolein (MP, Nu-Chek Prep) in 2:3 ratio (lipid:protein) to form a lipidic mesophase. The mesophase was homogenized in coupled syringes (Hamilton) by transferring the mesophase from one syringe to another until a homogeneous and gel-like material was formed(Nollert, 2004). 150 nl drops of a protein–mesophase mixture were spotted on a 96-well LCP glass sandwich plate (Marienfeld) and overlaid with 400 nL of precipitant solution by means of the NT8 crystallization robot (Formulatrix). The best crystals of OLPVR1 were obtained with a protein concentration of 20 mg/ml and 10 mM CaCl_2_, 10 mM MgCl_2_, 24% PEG 6000, 100mM Tris (pH 8.2) for MP lipid and 10 mM CaCl_2_, 10 mM MgCl_2_, 24% PEG 550, 100mM Tris (pH 8.2) for MO lipid (Hampton Research). The best crystals of O1O2 were obtained with a protein concentration of 20 mg/ml and 1.8 M Na_2_HPO_4_/KH_2_PO_4_ (pH 4.6). The crystals were grown at 22°C and appeared in 1 to 4 weeks. Once crystals reached their final size, crystallization wells were opened as described elsewhere (Li et al., 2012), and drops containing the protein-mesophase mixture were covered with 100 μl of the respective precipitant solution. For data collection harvested crystals were incubated for 5 min in the respective precipitant solutions.

#### Molecular dynamics

We used the refined 1.4 Å resolution crystallographic OLPVR1 structure for the initial conformation. All non-protein atoms except water were removed from the structure and the all-trans retinal molecule connected to the Lys were renamed using the retinol and retinal parameters for Charmm36 force field. The system then was prepared using Charmm GUI (Jo et al., 2008) input generator using the POPC lipid membrane and Tip3P water model. The resulting amount of lipids was 132, amount of water molecules - 8901, amount of sodium ions - 26, chlorine ions - 23, overall system size was 48350 atoms. Energy minimization and equilibration were performed in several steps with the gradual removal of spatial atomic constraints. The resulting simulation time was 1 μs (current time is 0.75μs). Simulations were performed using velocity-rescale thermostat at 303.15 K and Parrinello-Rahman semi isotropic barostat with Gromacs 2018.4 (Pronk et al., 2013).

## Acknowledgments

We thank C. Baeken for technical assistance. We acknowledge the Structural Biology Group of the European Synchrotron Radiation Facility (ESRF) for granting access to the synchrotron beamlines. We also acknowledge the Paul Scherrer Institut, Villigen, Switzerland for provision of synchrotron radiation beamtime at beamline X06SA of the SLS and would like to thank Dr. Anuschka Pauluhn for assistance. This work was supported by the common program of Agence Nationale de la Recherche (ANR), France and Deutsche Forschungsgemeinschaft (DFG), Germany (ANR-15-CE11-0029-02/FA 301/11-1), by the DFG Research Unit FOR 2518 (*DynIon*, project P4 to JPM, MA 7525/1-1), and by funding from Frankfurt: Cluster of Excellence Frankfurt Macromolecular Complexes by the Max Planck Society (to E.B.) and by the Commissariat à l’Energie Atomique et aux Energies Alternatives (Institut de Biologie Structurale)–Helmholtz-Gemeinschaft Deutscher Forschungszentren (Forschungszentrum Jülich) Special Terms and Conditions 5.1 specific agreement. This work used the platforms of the Grenoble Instruct-ERIC center (ISBG; UMS 3518 CNRS-CEA-UJF-EMBL) within the Grenoble Partnership for Structural Biology (PSB). Platform access was supported by FRISBI (ANR-10-INBS-05-02) and GRAL, a project of the University Grenoble Alpes graduate school (Ecoles Universitaires de Recherche) CBH-EUR-GS (ANR-17-EURE-0003). The reported study was funded by RFBR and CNRS according to the research project № 19-52-15017. DB was supported by grant ANR-14-CE09-0028. FRV was supported by grant VIREVO CGL2016-76273-P [AEI/FEDER, EU], (cofounded with FEDER funds). DZ and RA were supported by the Russian Foundation for Basic Research project number 17-00-00164.

## Author contributions

DZ, TB, DB and SV did molecular cloning, expressed and purified the proteins. DV, DS and IC measured the photocycle kinetics and analyzed the corresponding data. DZ and RA crystallized the proteins. KK and DZ harvested crystals, processed the data and solved the structures with the help of AP. AA and ASO performed electrophysiology experiments under EB and MV supervision. DZ, MR, TR and YA carried out the functional tests. GA and KS did the molecular dynamics experiments. RR and FRV performed metagenomic search and identified new viral rhodopsin sequences. DZ, NY and EK analyzed the viral rhodopsins sequences and their possible biological role. VG initiated and supervised the viral rhodopsins project. VG, GB, EK, MK and EB planned and guided the work. DZ, AA, KK and VG analyzed the data and prepared the manuscript with the input from all other authors.

## Supplemental text

### Unusual protein-lipid interaction in the OLPVR1 structure

Surprisingly, both OLPVR1 structures revealed a fragment of a lipid molecule (monopalmitolein, 97N) between the TM6 and TM7 helices immersed into a pore at the intracellular side of the protein and also a groove at the protein surface, which connects the pore and the cytoplasm, described earlier (Extended Data Figure 3a). The hydrophilic ‘head’ of the lipid is resolved nicely at high resolution and interacts directly with water molecules of the intracellular cavity (Figure 6b). We suspect that the pore is accidentally blocked by the lipid molecule of the host crystallization matrix. In order to address the potential structural importance of the lipid molecule and the stability of the OLPVR1 in its absence, we performed molecular dynamics simulations of OLPVR1 without the lipid fragment. During the simulation protein did not show any significant side chain displacement in the nearest vicinity of the lipid binding region (Extended Data Figure 3d). In addition, we designed a variant of the OLPVR1 with nine mutations (I169L, Y172G, F173A, V177L, F179A, V202F, I206F, Y207V, I211F) that we denote as O1O2 protein, which intracellular parts of TM6 and TM7 helices are supposed to mimic those of OLPVR2 from VR2 group (Yutin and Koonin, 2012) (Extended Data Figure 3c). We expressed, purified and crystallized O1O2 similarly to the wild type protein. O1O2 crystals diffracted up to 1.9Å and contained two protein molecules in the asymmetric unit, organized into the same dimer as demonstrated for the wild type OLPVR1 (Extended Data Figure 9). Interestingly, although the pore in the intracellular part of the O1O2 is similar to that of OLPVR1, we observed no lipid fragments in the lipid-binding region (Extended Data Figure 3b). Thus, lipid molecule most likely does not play a role in structural stabilization of the protein, but rather employ loose configuration between TM6 and TM7 helices to stuck inside.

### Comparison of the structures of group 1 viral rhodopsins

In order to underline similarities and discrepancies between two available structures of viral rhodopsins from group 1, we performed a detailed comparison of OLPVR1 (this study) and DTS rhodopsin of *Phaeocystis globosa* virus 12T (Needham et al., 2019) (PDB: 6J0O). The structures present nearly identical topology of the helices and superimpose with RMSD of 0.44 Å (Extended Data Figure 4). Similar to other viral rhodopsins from group 1, DTS rhodopsins conserve all residues that are predicted to be responsible for ion channeling activity. In particular, we observed nearly identical orientations of side chains of S11, E44, E51, N197 and E205 residues in both structures suggesting a similar configuration of those residues among all group 1viral channelrhodopsins. Interestingly, the DTS rhodopsin contains a lipid molecular fragment between helices TM6 and TM7 similar to those in OLPVR1 structure, supporting the hypothesis on the conservation of the pore structure between TM6 and TM7 in viral rhodopsins. Taken together, our data suggest that the lipid molecule should not be considered a functional part of the protein and it is rather an artifact of crystallization.

## Extended Data figures

**Extended Data Figure 1.**
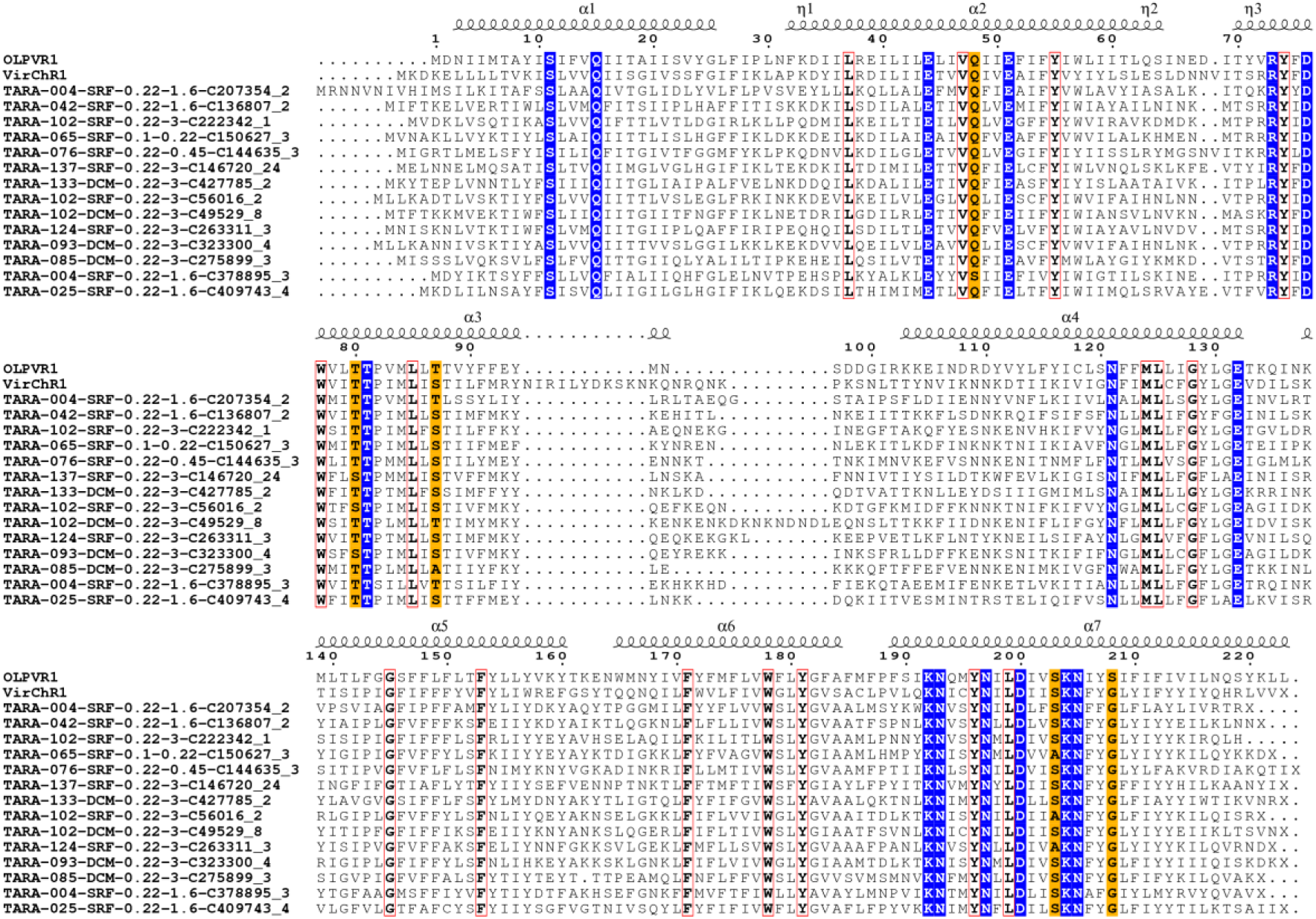
Sequence alignment of representative viral rhodopsins. The sequence alignment was created using UGENE software and ESPript3 online server. Secondary structure elements for OLPVR1 are shown as coils. Fully conservative residues are indicated with blue color and red frames for charged and non-charged amino acids respectively. Highly conservative residues forming ion channeling constrictions are additionally colored orange.

**Extended Data Figure 2.**
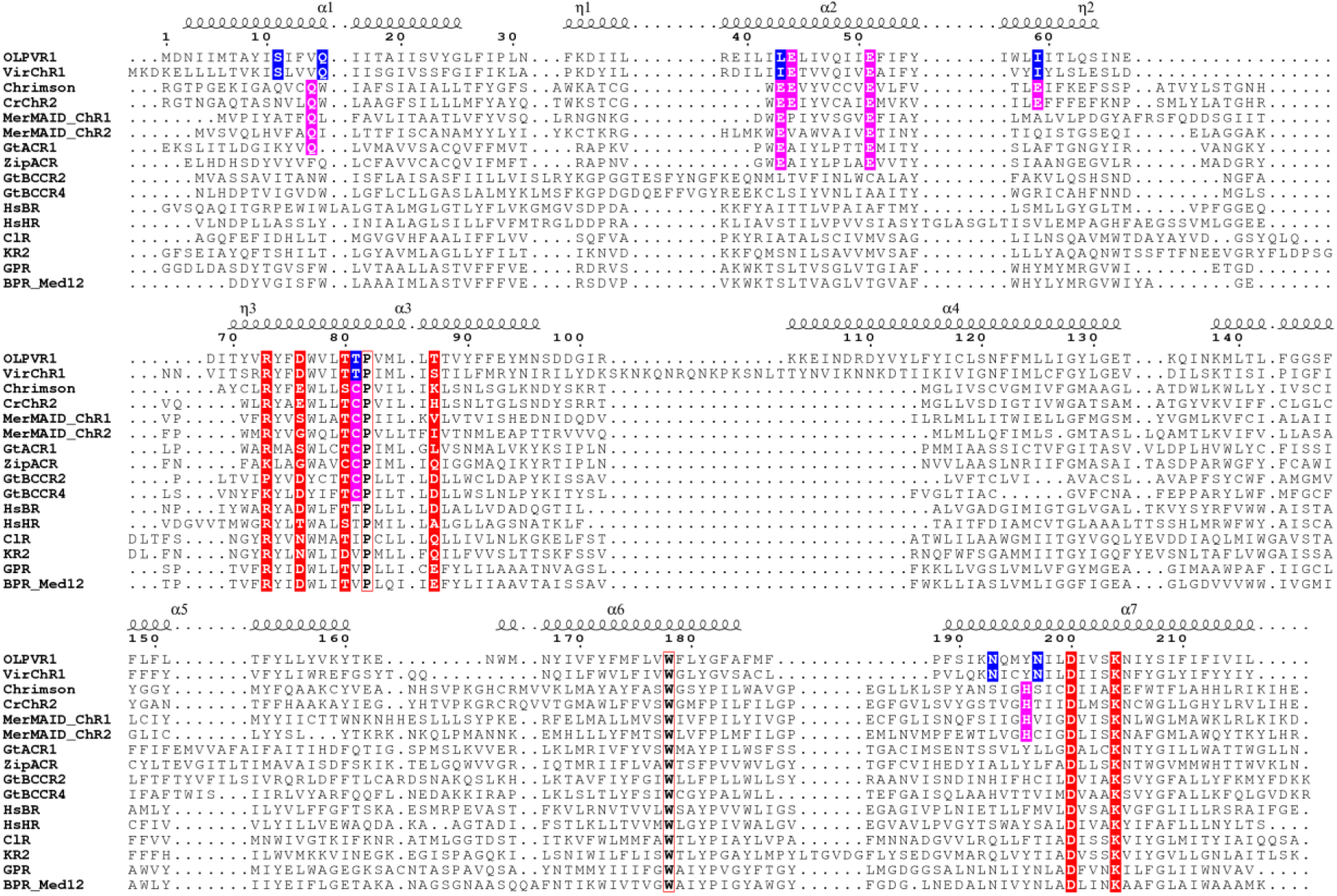
Sequence alignment of microbial opsin genes. The sequences with their Genbank IDs in parenthesis are OLPVR1 (ADX06642.1), VirChR1 (TARA-146-SRF-0.22-3-C376786_1, this study), Chrimson (AHH02126.1), *Cr*ChR2 (ABO64386.1), MerMAID_ChR1(Oppermann et al., 2019), MerMAID_ChR2(Oppermann et al., 2019), GtACR1 (AKN63094.1), ZipACR (APZ76709.1), GtBCCR2 (ANC73518.1), GtBCCR4 (ARQ20888.1), *Hs*BR (WP_010903069.1), HsHR (P15647.1), ClR (PDB ID: 5G28), KR2 (BAN14808.1), GPR (AAK30191.1), BPR_Med12 (PDB ID: 4JQ6). Residues included in 6-letter motif are colored red, fully conservative residues are indicated with red frames. Key amino acids involved in ion channeling are colored blue and purple for viral rhodopsin group unique positions and channelrhodopsin conservative positions respectively.

**Extended Data Figure 3.**
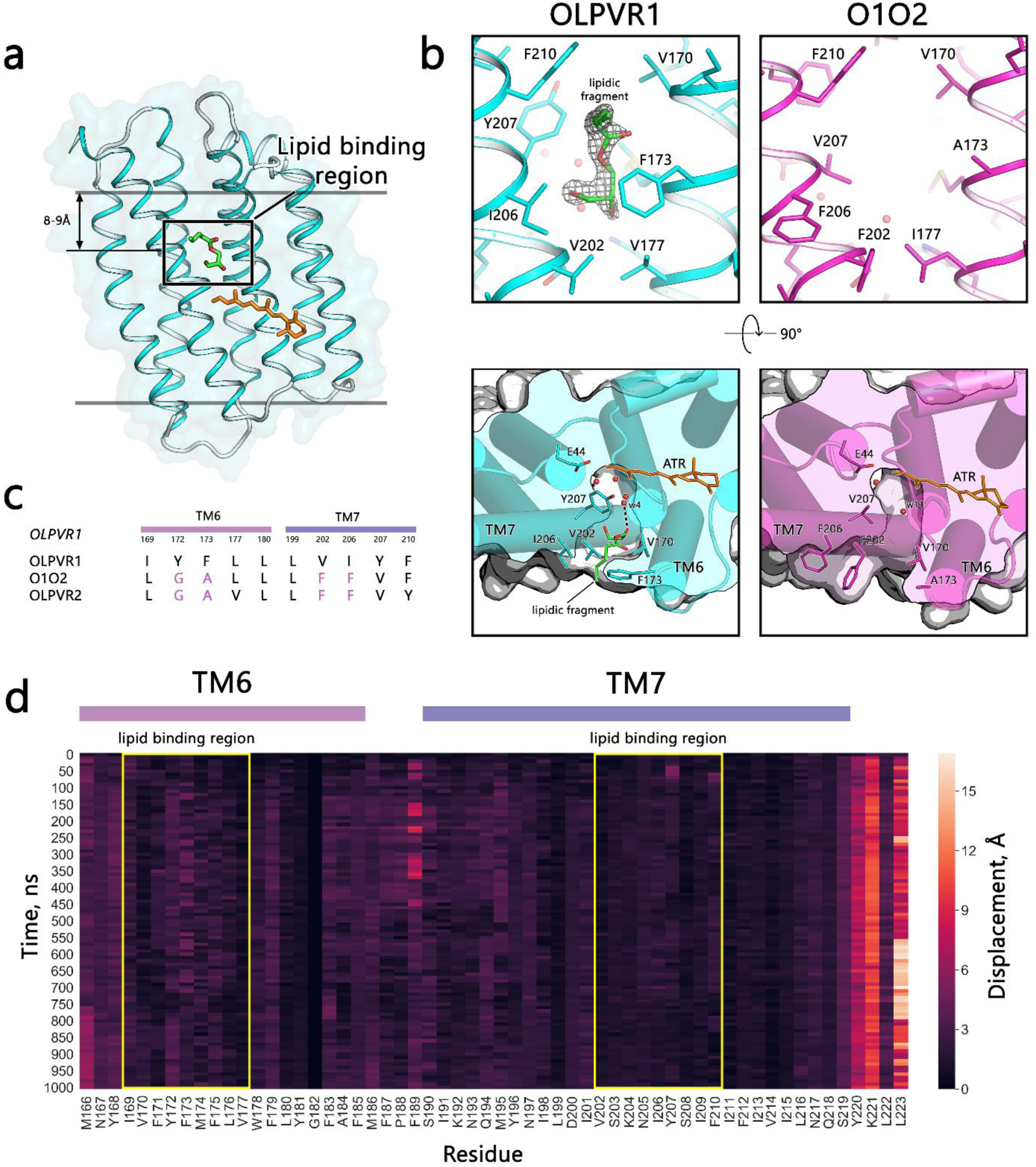
Unusual OLPVR1-lipid interaction. (a) Lipid binding region (LBR) in OLPVR1 structure between TM6 and TM7 helices. The lipidic fragment is drawn as green sticks. (b) The magnified view of protein-lipid interaction region in OLPVR1 (left) and O1O2 (right). 2F_o_ – F_c_ map of the lipidic fragment between TM6 and TM7 helices is contoured at 1.5σ and shown as gray mesh. No lipidic fragment in the LBR in O1O2 structure was observed. (c) Sequence alignment of the TM6 and TM7 helices of OLPVR1, OLPVR2(Bratanov et al., 2019) and O1O2 mutant. Major mutations implemented are colored purple. (d) Heat plot of side chain displacement (root mean square deviation, RMSD) of residues from TM6 and TM7 at 1 μs scale after lipidic fragment (97N 302 molecule) removal.

**Extended Data Figure 4.**
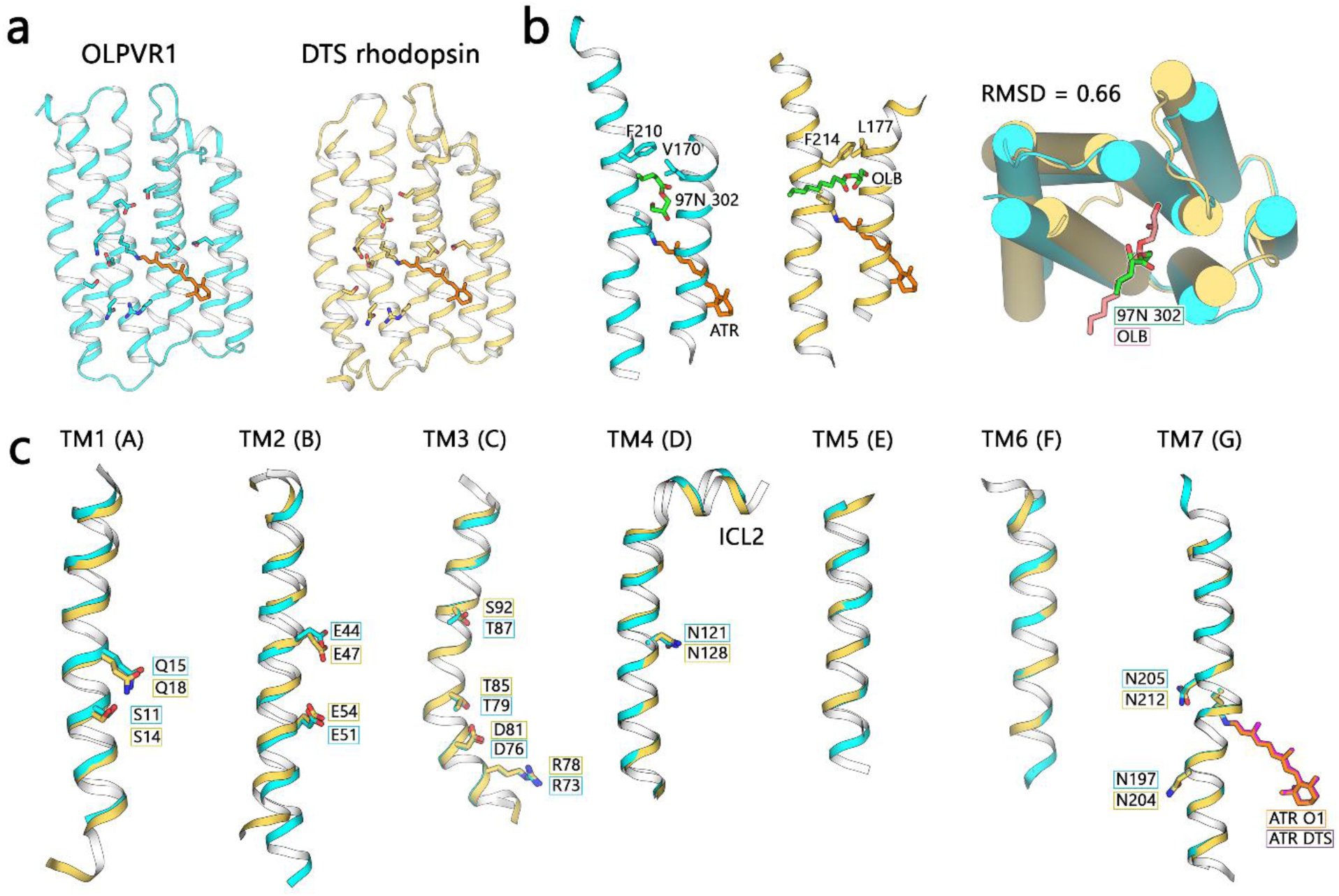
Extended comparison of OLPVR1 and DTS rhodopsin structures. (a) Crystal structures of OLPVR1 and DTS rhodopsins (PDB: 6JO0). (b) Extended overview of the lipid molecule fragment orientation between TM6 and TM7 helices in viral rhodopsin structures viewed parallel to membrane (left) and from the intracellular side (right). (c) Individual TM helices are shown after superimposition of the OLPVR1 and DTS rhodopsin. Important residues are additionally shown as sticks.

**Extended Data Figure 5.**
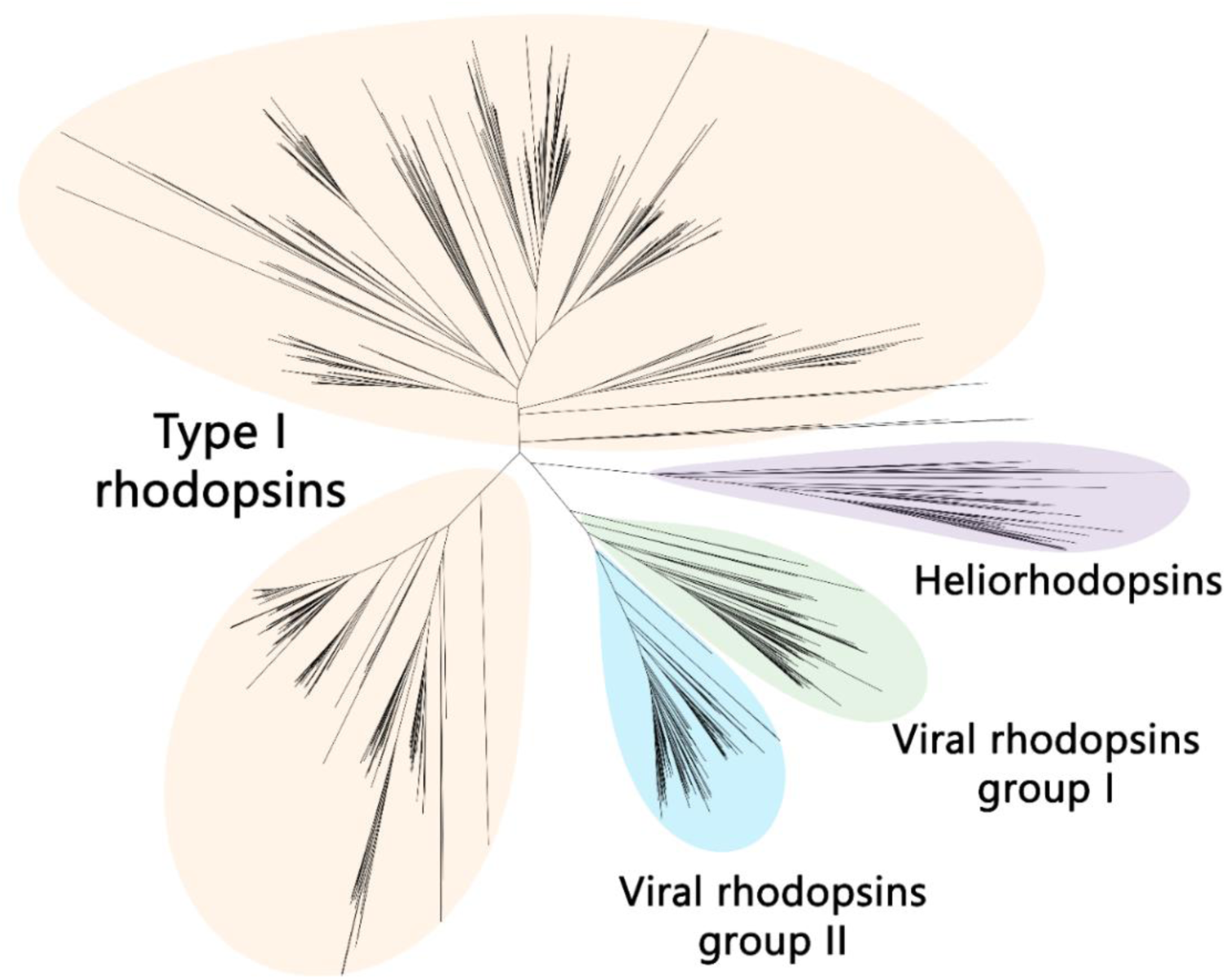
Unrooted phylogenetic tree of known microbial rhodopsins. Viral rhodopsins (group I and II), heliorhodopsins (Pushkarev et al., 2018) and other microbial rhodopsins (type-I rhodopsins) groups are indicated with color. Analyzed base of microbial rhodopsins consists of proteins retrieved from TARA project and proteins predicted with InterPro software(Mitchell et al., 2019). Phylogenetic tree was created using UGENE and iTOL software (Letunic and Bork, 2016).

**Extended Data Figure 6.**
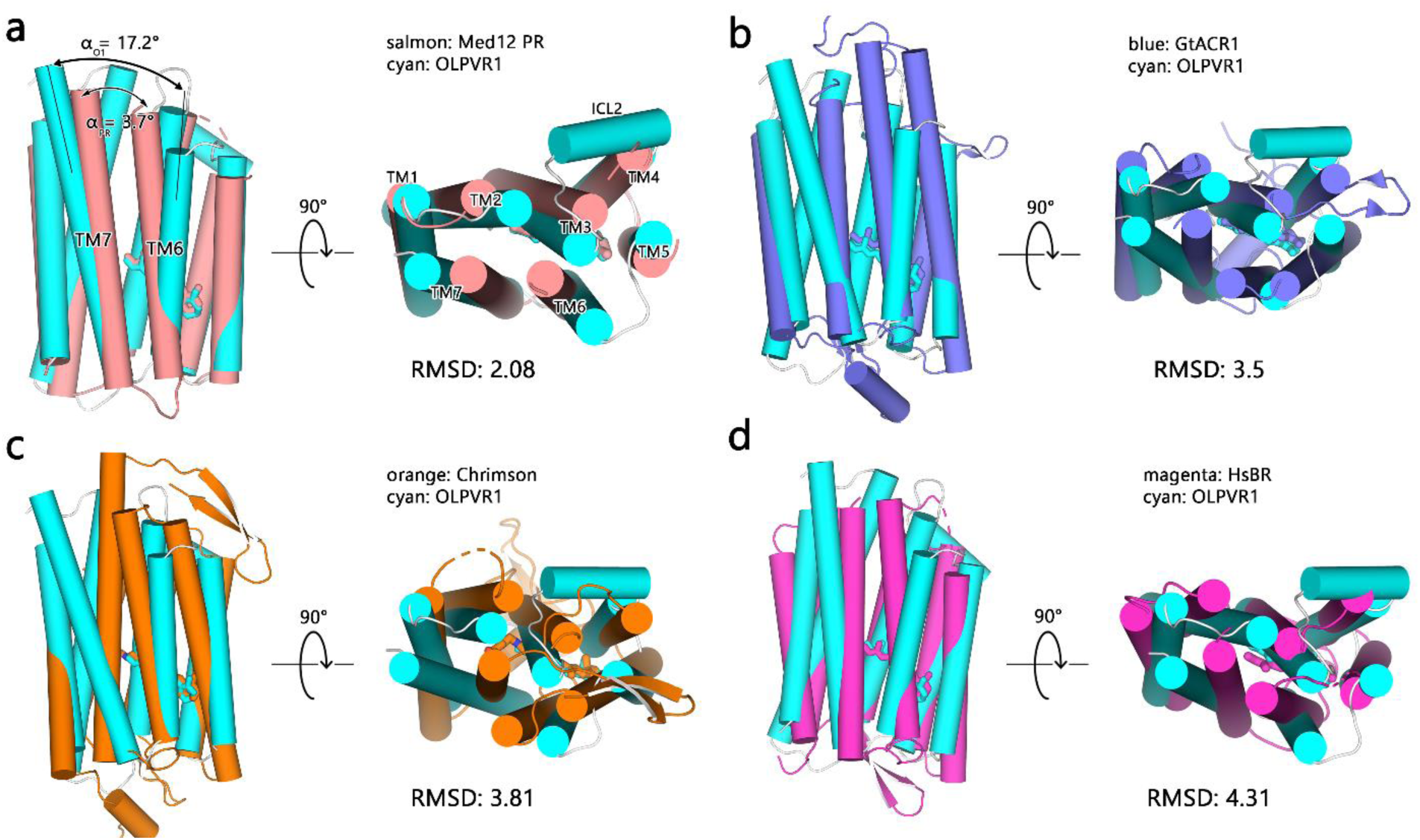
Structure alignment of OLPVR1 with other rhodopsins. Side views and views from extracellular side of pair structure alignments of OLPVR1 with (a) Med12 PR (PDB ID: 4JQ6), (b) GtACR1 (PDB: 6CSN), (c) Chrimson (PDB: 5ZIH) and (d) *Hs*BR (PDB: 1C3W). Root mean square deviations (RMSD) of the alignments are additionally indicated.

**Extended Data Figure 7.**
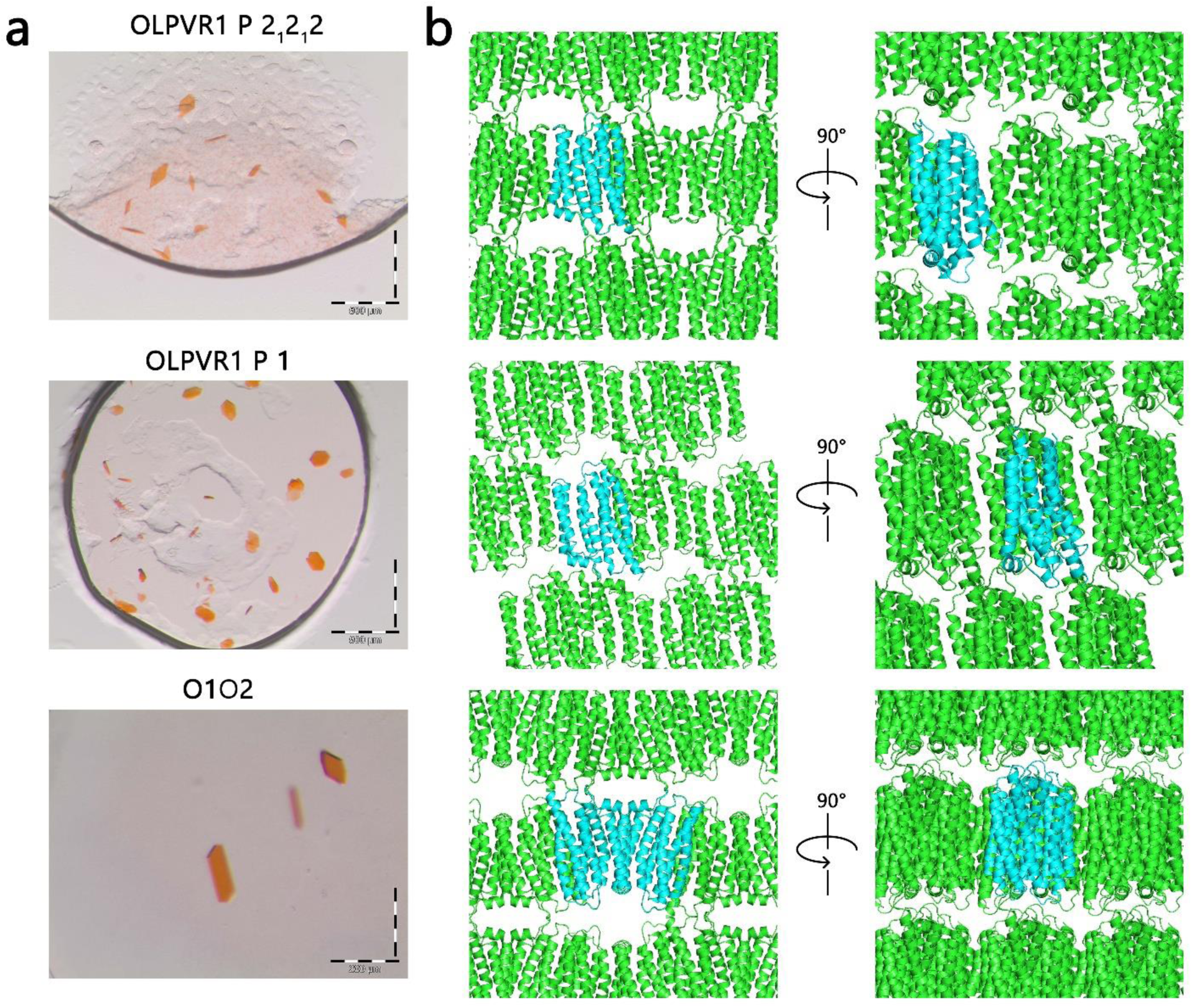
Crystal packing of OLPVR1 protein. (a) Magnified images of the OLPVR1 and O1O2 crystals used for structure determination. (b) Lattice packing of the crystals, viewed in two principal orientations. Asymmetric unit protein content is colored cyan for each structure.

**Extended Data Figure 8.**
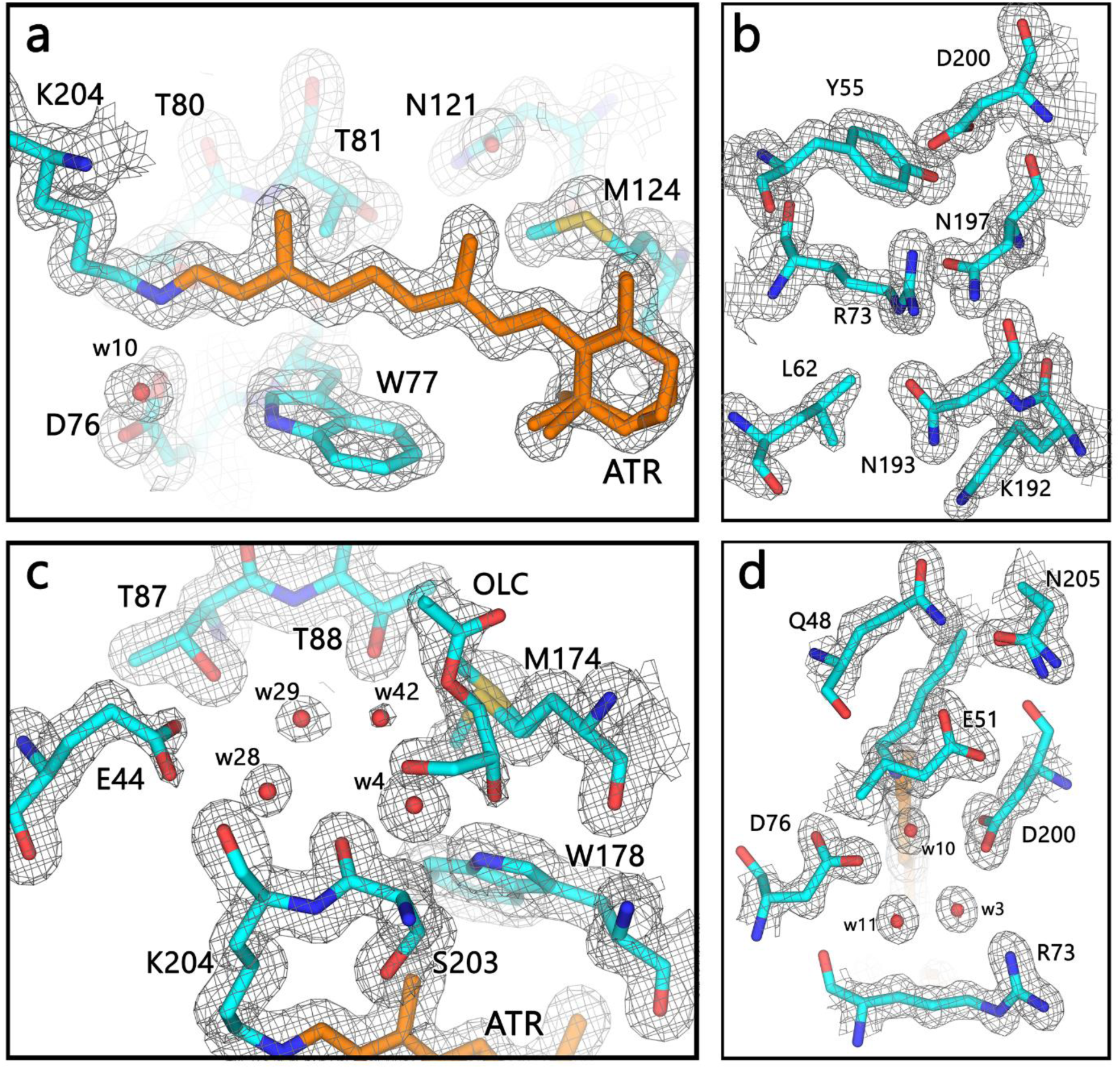
Examples of electron density map of OLPVR1 structure. 2F_o_ – F_c_ maps (gray mesh, contoured at 1.5σ) for the (a) retinal pocket, (b) intracellular gate, (c) extracellular gate and (d) central gate. The water molecules and retinal cofactor in 2 conformations are clearly resolved in the structure. 1.4Å resolution dataset was collected with the single crystals.

**Extended Data Figure 9.**
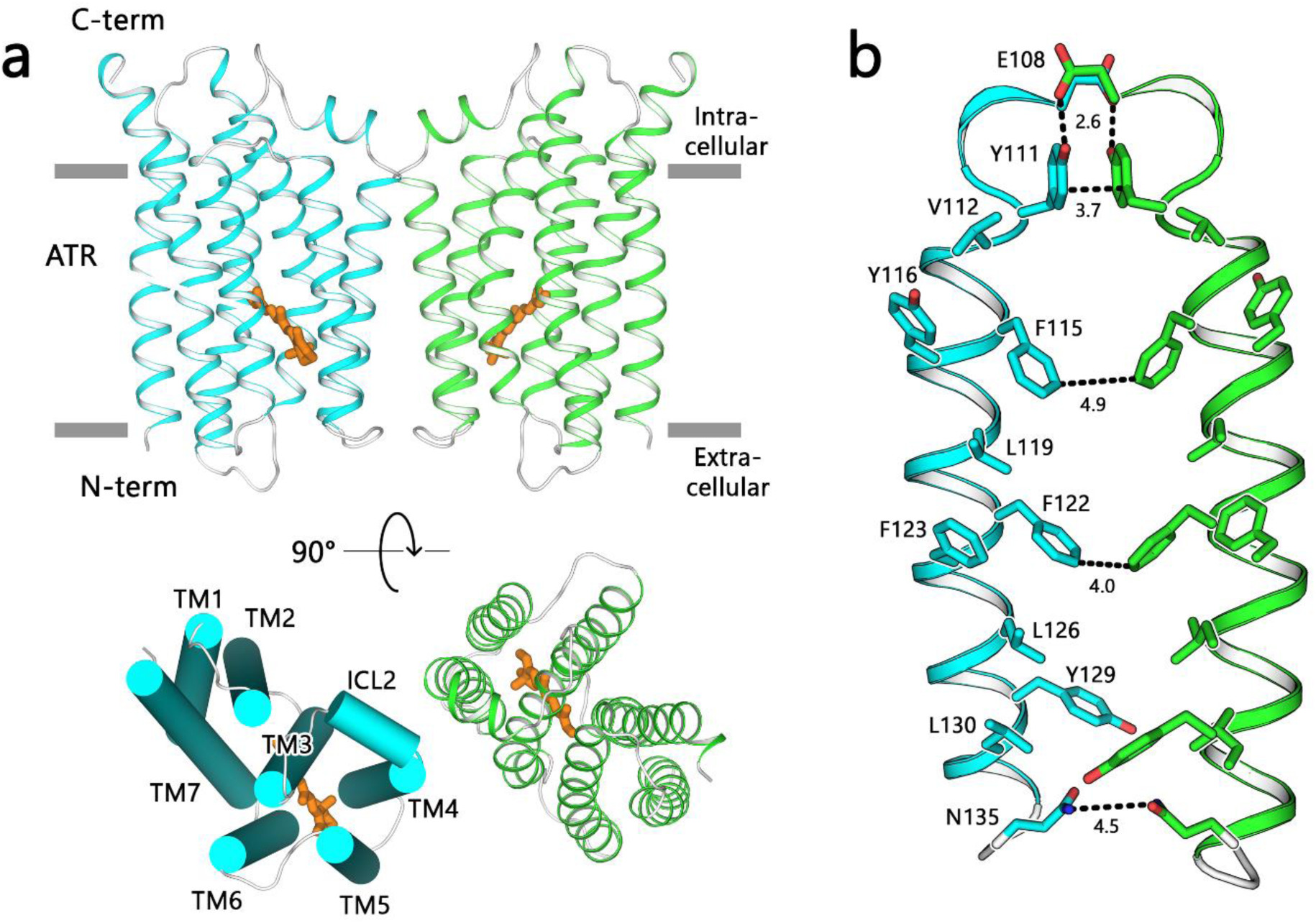
Putative dimer organization of OLPVR1. (a) Side view and view from extracellular side of the OLPVR1 putative dimer, observed for P2_1_2_1_2 structure. (b) Putative dimerization interface of OLPVR1. Distances between residues involved in dimer formation are drawn with dashed lines. Residues involved into dimer formation are named in one of the protomers.

**Extended Data Figure 10.**
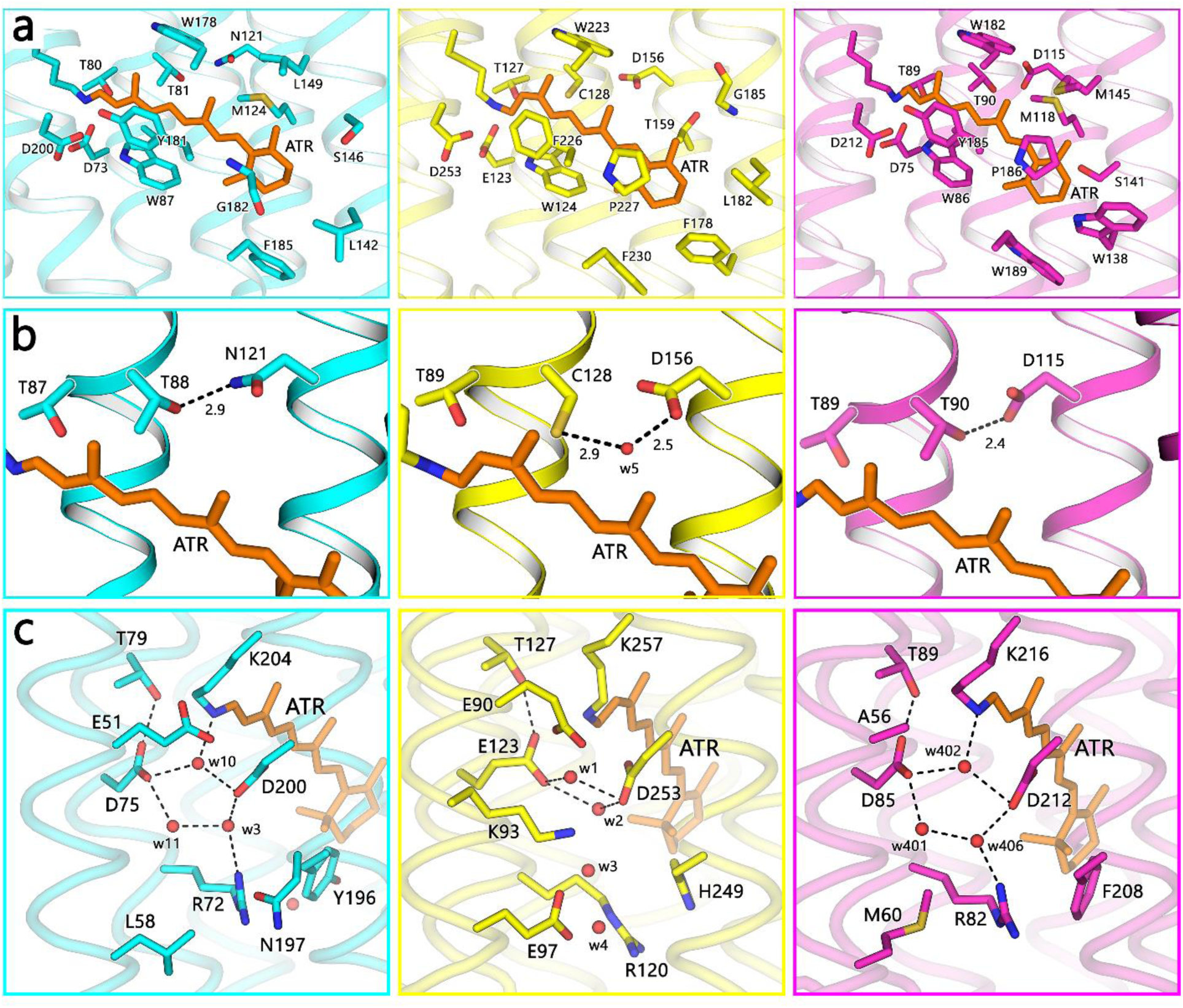
Retinal pocket and DC pair region architecture. (a) Retinal pockets, (b) DC pair regions and (c) Schiff base regions of OLPVR1 (left), *Cr*ChR2 (middle) and *Hs*BR (right) are shown in cyan, yellow and purple colors respectively. All-*trans* retinal cofactor is depicted orange. Important residues are numbered and depicted as sticks. Salt bridge interactions are shown with dashed lines. Distances between DC-pair partnering residues are indicated in angstroms.

**Extended Data Figure 11.**
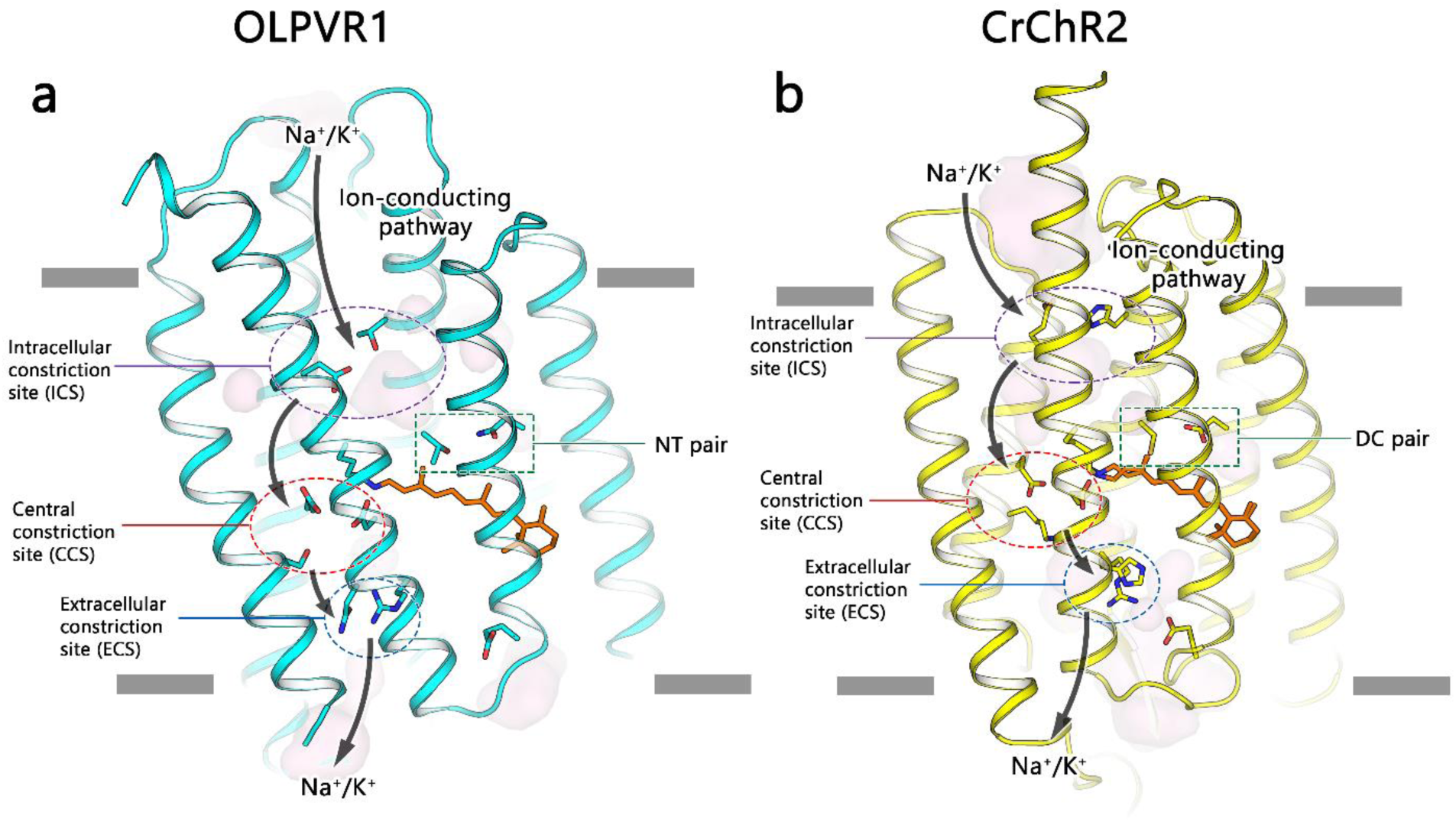
Proposed ion-conducting pathway of OLPVR1. Three consecutive constriction sites and proposed cation pathway of (a) OLPVR1 and (b) *Cr*ChR2 proteins. The proposed ion pathway is shown only in one direction for clarity. Important residues and depicted as sticks. All-*trans* retinal is depicted with orange color. NT and DC pairs are additionally indicated.

**Extended Data Figure 12.**
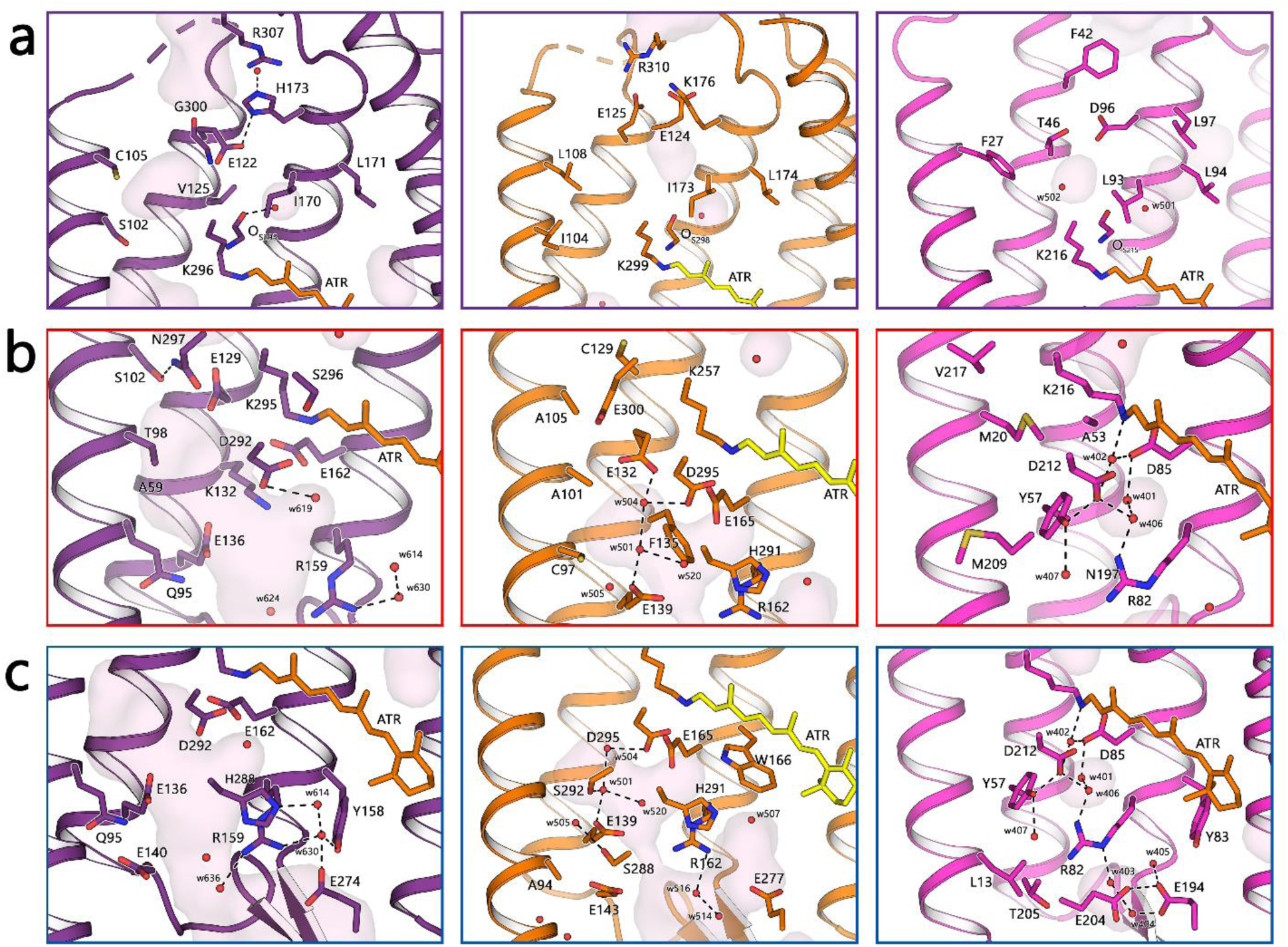
Organization of ion pathway constriction sites (gates) in other microbial rhodopsins. Magnified view of the (a) intracellular constriction site (gate), (b) central constriction site (gate) and (c) extracellular constriction site (gate) regions in C1C2 (left, PDB code: 3UG9), Chrimson (middle, PDB code: 5ZIH) and *Hs*BR (right, PDB code: 1C3W) structures, colored indigo, orange and magenta respectively. Water accessible cavities were calculated using HOLLOW and are presented as pink surfaces(Ho and Gruswitz, 2008).

**Extended Data Table 1.** Crystallographic data collection and refinement statistics.

| **Data collection** | OLPVR1_P1 | OLPVR1_P21212 | O1O2 |
| --- | --- | --- | --- |
| Space group | P 1 | P 2<sub>1</sub> 2<sub>1</sub> 2 | I 1 2 1 |
| Cell dimensions |  |  |  |
| *a*, *b*, *c* (Å) | 40.18, 56.97, 62.34 | 46.53, 115.80, 53.53 | 44.40, 103.40, 111.38 |
| $\alpha$, $\beta$, $\gamma$ (°) | 113.90, 90.01, 91.49 | 90, 90, 90 | 90, 90.32, 90 |
| Wavelength (Å) | 0.976 | 0.976 | 0.978 |
| Resolution (Å) | 40.16-1.6 (1.63-1.60) | 48.59-1.4 (1.42-1.40) | 41.33-1.96 (2.00-1.96) |
| *R*<sub>merge</sub> (%) | 5.8 (116.1) | 7.2 (181.0) | 5.1 (119.1) |
| *I*/σ*I* | 6.1 (0.7) | 12.3 (1.0) | 12.2 (0.9) |
| *CC*<sub>1/2</sub> (%) | 99.8 (53.3) | 99.8 (45.3) | 99.9 (50.8) |
| Completeness (%) | 95.2 (94.2) | 100.0 (99.9) | 99.5 (95.4) |
| Unique reflections | 63,486 (3098) | 57,845 (2820) | 36,098 (2448) |
| Multiplicity | 3.5 (3.6) | 6.3 (6.3) | 3.5 (3.4) |
| **Refinement** |  |  |  |
| Resolution (Å) | 20-1.60 | 20-1.40 | 20-1.96 |
| No. reflections | 60,315 | 54,864 | 34,293 |
| *R*<sub>work</sub>/*R*<sub>free</sub> (%) | 18.8/21.3 | 14.2/18.6 | 20.2/24.6 |
| No. atoms |  |  |  |
| Protein | 3915 | 2012 | 3776 |
| Retinal | 40 | 40 | 40 |
| Water | 98 | 107 | 52 |
| Lipids | 289 | 220 | 32 |
| B-factors (Å<sup>2</sup>) |  |  |  |
| Protein | 31 | 24 | 45 |
| Retinal | 23 | 17 | 35 |
| Water | 36 | 37 | 42 |
| Lipids | 54 | 46 | 58 |
| R.m.s. deviations |  |  |  |
| Bond lengths (Å) | 0.0030 | 0.0041 | 0.0021 |
| Bond angles (°) | 1.0662 | 0.8917 | 1.0525 |

## Bibliography

Béjà, O., Aravind, L., Koonin, E. V, Suzuki, M.T., Hadd, A., Nguyen, L.P., Jovanovich, S.B., Gates, C.M., Feldman, R.A., Spudich, J.L., et al. (2000). Bacterial Rhodopsin : Evidence for a New Type of Phototrophy in the Sea. Science 289, 1902–1906.

Berndt, A., and Deisseroth, K. (2015). Expanding the optogenetics toolkit. Science 349, 590–591.

Berndt, A., Yizhar, O., Gunaydin, L.A., Hegemann, P., and Deisseroth, K. (2009). Bi-stable neural state switches. Nat. Neurosci. 12, 229–234.

Berndt, A., Lee, S.Y., Ramakrishnan, C., and Deisseroth, K. (2014). Structure-Guided Transformation of Channelrhodopsin into a Light-Activated Chloride Channel. Science 344, 420–424.

Böhm, M., Boness, D., Fantisch, E., Erhard, H., Frauenholz, J., Kowalzyk, Z., Marcinkowski, N., Kateriya, S., Hegemann, P., and Kreimer, G. (2019). Channelrhodopsin-1 Phosphorylation Changes with Phototactic Behavior and Responds to Physiological Stimuli in *Chlamydomonas*. Plant Cell 31, 886–910.

Bratanov, D., Kovalev, K., Machtens, J.-P., Astashkin, R., Chizhov, I., Soloviov, D., Volkov, D., Polovinkin, V., Zabelskii, D., Mager, T., et al. (2019). Unique structure and function of viral rhodopsins. Nat. Commun. 10, 4939.

Buchfink, B., Xie, C., and Huson, D.H. (2015). Fast and sensitive protein alignment using DIAMOND. Nat. Methods 12, 59–60.

Caffrey, M., and Cherezov, V. (2009). Crystallizing membrane proteins using lipidic mesophases. Nat. Protoc. 4, 706–731.

Charlson, R.J., Lovelock, J.E., Andreae, M.O., and Warren, S.G. (1987). Oceanic phytoplankton, atmospheric sulphur, cloud albedo and climate. Nature 326, 655–661.

Chizhov, I., Chernavskii, D.S., Engelhard, M., Mueller, K.H., Zubov, B. V., and Hess, B. (1996). Spectrally silent transitions in the bacteriorhodopsin photocycle. Biophys. J. 71, 2329–2345.

Crooks, G.E. (2004). WebLogo: A Sequence Logo Generator. Genome Res. 14, 1188–1190.

Deisseroth, K. (2015). Optogenetics: 10 years of microbial opsins in neuroscience. 18, 1213–1225.

Ernst, O.P., Lodowski, D.T., Elstner, M., Hegemann, P., Brown, L.S., and Kandori, H. (2014). Microbial and animal rhodopsins: Structures, functions, and molecular mechanisms. Chem. Rev. 114, 126–163.

Fenalti, G., Giguere, P.M., Katritch, V., Huang, X.-P., Thompson, A.A., Cherezov, V., Roth, B.L., and Stevens, R.C. (2014). Molecular control of δ-opioid receptor signalling. Nature 506, 191–196.

Filée, J. (2018). Giant viruses and their mobile genetic elements: the molecular symbiosis hypothesis. Curr. Opin. Virol. 33, 81–88.

Freier, E., Wolf, S., and Gerwert, K. (2011). Proton transfer via a transient linear water-molecule chain in a membrane protein.

Gómez-Consarnau, L., Raven, J.A., Levine, N.M., Cutter, L.S., Wang, D., Seegers, B., Arístegui, J., Fuhrman, J.A., Gasol, J.M., and Sañudo-Wilhelmy, S.A. (2019). Microbial rhodopsins are major contributors to the solar energy captured in the sea. Sci. Adv. 5, eaaw8855.

Gordeliy, V.I., Labahn, J., Moukhametzianov, R., Efremov, R., Granzin, J., Schlesinger, R., Büldt, G., Savopol, T., Scheidig, A.J., Klare, J.P., et al. (2002). Molecular basis of transmembrane signalling by sensory rhodopsin II-transducer complex. Nature 419, 484–487.

Govorunova, E.G., Sineshchekov, O.A., Janz, R., Liu, X., and Spudich, J.L. (2015). Natural light-gated anion channels: A family of microbial rhodopsins for advanced optogenetics. Science.

Govorunova, E.G., Sineshchekov, O.A., and Spudich, J.L. (2016). Structurally Distinct Cation Channelrhodopsins from Cryptophyte Algae. Biophys. J. 110, 2302–2304.

Govorunova, E.G., Sineshchekov, O.A., Rodarte, E.M., Janz, R., Morelle, O., Melkonian, M., Wong, G.K., and Spudich, J.L. (2017). The Expanding Family of Natural Anion Channelrhodopsins Reveals Large Variations in Kinetics, Conductance, and Spectral Sensitivity. Nat. Publ. Group 1–10.

Gradinaru, V., Zhang, F., Ramakrishnan, C., Mattis, J., Prakash, R., Diester, I., Goshen, I., Thompson, K.R., and Deisseroth, K. (2010). Molecular and Cellular Approaches for Diversifying and Extending Optogenetics. Cell 141, 154–165.

Gushchin, I., and Gordeliy, V. (2018). Microbial rhodopsins.

Gushchin, I., Chervakov, P., Kuzmichev, P., Popov, A.N., Round, E., Borshchevskiy, V., Ishchenko, A., Petrovskaya, L., Chupin, V., Dolgikh, D.A., et al. (2013). Structural insights into the proton pumping by unusual proteorhodopsin from nonmarine bacteria. Proc. Natl. Acad. Sci. 110, 12631–12636.

Gushchin, I., Shevchenko, V., Polovinkin, V., Kovalev, K., Alekseev, A., Round, E., Borshchevskiy, V., Balandin, T., Popov, A., Gensch, T., et al. (2015). Crystal structure of a light-driven sodium pump. Nat. Struct. Mol. Biol. 22, 390–396.

Harz, H., and Hegemann, P. (1991). Rhodopsin-regulated calcium currents in Chlamydomonas. Nature 351, 489– 491.

Ho, B.K., and Gruswitz, F. (2008). HOLLOW: Generating accurate representations of channel and interior surfaces in molecular structures. BMC Struct. Biol.

Hyatt, D., Chen, G.L., LoCascio, P.F., Land, M.L., Larimer, F.W., and Hauser, L.J. (2010). Prodigal: Prokaryotic gene recognition and translation initiation site identification. BMC Bioinformatics 11, 119.

Jékely, G. (2009). Evolution of phototaxis. Philos. Trans. R. Soc. B Biol. Sci. 364, 2795–2808.

Jo, S., Kim, T., Iyer, V.G., and Im, W. (2008). CHARMM-GUI: A web-based graphical user interface for CHARMM. J. Comput. Chem. 29, 1859–1865.

Jorgensen, B.B., Cohen, Y., and Des Marais, D.J. (1987). Photosynthetic action spectra and adaptation to spectral light distribution in a benthic cyanobacterial mat. Appl. Environ. Microbiol. 53, 879.

Käll, L., Krogh, A., and Sonnhammer, E.L.L. (2004). A combined transmembrane topology and signal peptide prediction method. J. Mol. Biol. 338, 1027–1036.

Kato, H.E., Zhang, F., Yizhar, O., Ramakrishnan, C., Nishizawa, T., Hirata, K., Ito, J., Aita, Y., Tsukazaki, T., Hayashi, S., et al. (2012). Crystal structure of the channelrhodopsin light-gated cation channel. Nature 482, 369– 374.

Kato, H.E., Inoue, K., Abe-Yoshizumi, R., Kato, Y., Ono, H., Konno, M., Hososhima, S., Ishizuka, T., Hoque, M.R., Kunitomo, H., et al. (2015). Structural basis for Na + transport mechanism by a light-driven Na + pump. Nature 521, 48–53.

Kato, H.E., Kim, Y.S., Paggi, J.M., Evans, K.E., Allen, W.E., Richardson, C., Inoue, K., Ito, S., Ramakrishnan, C., Fenno, L.E., et al. (2018). Structural mechanisms of selectivity and gating in anion channelrhodopsins. Nature 561, 349–354.

Kianianmomeni, A., Stehfest, K., Nematollahi, G., Hegemann, P., and Hallmann, A. (2009). Channelrhodopsins of *Volvox carteri* Are Photochromic Proteins That Are Specifically Expressed in Somatic Cells under Control of Light, Temperature, and the Sex Inducer. Plant Physiol. 151, 347–366.

Kim, Y.S., Kato, H.E., Yamashita, K., Ito, S., Inoue, K., Ramakrishnan, C., Fenno, L.E., Evans, K.E., Paggi, J.M., Dror, R.O., et al. (2018). Crystal structure of the natural anion-conducting channelrhodopsin GtACR1. Nature 561, 343–348.

Klapoetke, N.C., Murata, Y., Kim, S.S., Pulver, S.R., Birdsey-Benson, A., Cho, Y.K., Morimoto, T.K., Chuong, A.S., Carpenter, E.J., Tian, Z., et al. (2014). Independent optical excitation of distinct neural populations. Nat. Methods 11, 338–346.

Kovalev, K., Polovinkin, V., Gushchin, I., Alekseev, A., Shevchenko, V., Borshchevskiy, V., Astashkin, R., Balandin, T., Bratanov, D., Vaganova, S., et al. (2019). Structure and mechanisms of sodium-pumping KR2 rhodopsin. Sci. Adv. 5, eaav2671.

Kuhne, J., Vierock, J., Tennigkeit, S.A., Dreier, M., Wietek, J., Petersen, D., Gavriljuk, K., El-Mashtoly, S.F., Hegemann, P., and Gerwert, K. (2019). Unifying photocycle model for light adaptation and temporal evolution of cation conductance in channelrhodopsin-2. Proc. Natl. Acad. Sci. 201818707.

Letunic, I., and Bork, P. (2016). Interactive tree of life (iTOL) v3: an online tool for the display and annotation of phylogenetic and other trees. Nucleic Acids Res. 44, W242–W245.

Li, W., and Godzik, A. (2006). Cd-hit: a fast program for clustering and comparing large sets of protein or nucleotide sequences. Bioinforma. Oxf. Engl. 22, 1658–1659.

Li, D., Boland, C., Aragao, D., Walsh, K., and Caffrey, M. (2012). Harvesting and Cryo-cooling Crystals of Membrane Proteins Grown in Lipidic Mesophases for Structure Determination by Macromolecular Crystallography. J. Vis. Exp. 1–7.

Lin, J.Y. (2011). A user’s guide to channelrhodopsin variants: features, limitations and future developments: A user’s guide to channelrhodopsin variants. Exp. Physiol. 96, 19–25.

Lin, J.Y., Sann, S.B., Zhou, K., Nabavi, S., Proulx, C.D., Malinow, R., Jin, Y., and Tsien, R.Y. (2013). Optogenetic Inhibition of Synaptic Release with Chromophore-Assisted Light Inactivation (CALI). Neuron 79, 241–253.

Lindell, D., Jaffe, J.D., Johnson, Z.I., Church, G.M., and Chisholm, S.W. (2005). Photosynthesis genes in marine viruses yield proteins during host infection. Nature 438, 86–89.

Lomize, M.A., Pogozheva, I.D., Joo, H., Mosberg, H.I., and Lomize, A.L. (2012). OPM database and PPM web server: Resources for positioning of proteins in membranes. Nucleic Acids Res.

López, J.L., Golemba, M., Hernández, E., Lozada, M., Dionisi, H., Jansson, J.K., Carroll, J., Lundgren, L., Sjöling, S., and Cormack, W.P.M. (2017). Microbial and viral-like rhodopsins present in coastal marine sediments from four polar and subpolar regions. FEMS Microbiol. Ecol. 93, 1–9.

Luecke, H., Schobert, B., Richter, H.-T., Cartailler, J.-P., and Lanyi, J.K. (1999). Structure of bacteriorhodopsin at 1.55 Å resolution 11Edited by D. C. Rees. J. Mol. Biol. 291, 899–911.

Mitchell, A.L., Attwood, T.K., Babbitt, P.C., Blum, M., Bork, P., Bridge, A., Brown, S.D., Chang, H.-Y., El-Gebali, S., Fraser, M.I., et al. (2019). InterPro in 2019: improving coverage, classification and access to protein sequence annotations. Nucleic Acids Res. 47, D351–D360.

Moore, S.K., Trainer, V.L., Mantua, N.J., Parker, M.S., Laws, E.A., Backer, L.C., and Fleming, L.E. (2008). Impacts of climate variability and future climate change on harmful algal blooms and human health. Environ. Health 7, S4.

Mukherjee, S., Hegemann, P., and Broser, M. (2019). Enzymerhodopsins: novel photoregulated catalysts for optogenetics. Curr. Opin. Struct. Biol. 57, 118–126.

Nack, M., Radu, I., Gossing, M., Bamann, C., Bamberg, E., von Mollard, G.F., and Heberle, J. (2010). The DC gate in Channelrhodopsin-2: crucial hydrogen bonding interaction between C128 and D156. Photochem. Photobiol. Sci. 9, 194.

Nagel, G., Szellas, T., Huhn, W., Kateriya, S., Adeishvili, N., Berthold, P., Ollig, D., Hegemann, P., and Bamberg, E. (2003). Channelrhodopsin-2, a directly light-gated cation-selective membrane channel. Proc. Natl. Acad. Sci. 100, 13940–13945.

Needham, D.M., Yoshizawa, S., Hosaka, T., Poirier, C., Choi, C.J., Hehenberger, E., Irwin, N.A.T., Wilken, S., Yung, C.-M., Bachy, C., et al. (2019). A distinct lineage of giant viruses brings a rhodopsin photosystem to unicellular marine predators. Proc. Natl. Acad. Sci. 116, 20574–20583.

Nollert, P. (2004). Lipidic cubic phases as matrices for membrane protein crystallization. Methods 34, 348–353.

Nultsch, W., Pfau, J., and Dolle, R. (1986). Effects of calcium channel blockers on phototaxis and motility of Chlamydomonas reinhardtii. Arch. Microbiol. 144, 393–397.

Oda, K., Vierock, J., Oishi, S., Rodriguez-Rozada, S., Taniguchi, R., Yamashita, K., Wiegert, J.S., Nishizawa, T., Hegemann, P., and Nureki, O. (2018). Crystal structure of the red light-activated channelrhodopsin Chrimson. Nat. Commun. 9, 3949.

Okada, T., Sugihara, M., Bondar, A.-N., Elstner, M., Entel, P., and Buss, V. (2004). The Retinal Conformation and its Environment in Rhodopsin in Light of a New 2.2Å Crystal Structure. J. Mol. Biol. 342, 571–583.

Okonechnikov, K., Golosova, O., and Fursov, M. (2012). Unipro UGENE: a unified bioinformatics toolkit. Bioinformatics 28, 1166–1167.

Oppermann, J., Fischer, P., Silapetere, A., Liepe, B., Rodriguez-Rozada, S., Flores-Uribe, J., Peter, E., Keidel, A., Vierock, J., Kaufmann, J., et al. (2019). MerMAIDs: a family of metagenomically discovered marine anion-conducting and intensely desensitizing channelrhodopsins. Nat. Commun. 10, 3315.

Price, M.N., Dehal, P.S., and Arkin, A.P. (2010). FastTree 2 - Approximately maximum-likelihood trees for large alignments. PLoS ONE 5, e9490.

Pronk, S., Páll, S., Schulz, R., Larsson, P., Bjelkmar, P., Apostolov, R., Shirts, M.R., Smith, J.C., Kasson, P.M., van der Spoel, D., et al. (2013). GROMACS 4.5: a high-throughput and highly parallel open source molecular simulation toolkit. Bioinformatics 29, 845–854.

Pushkarev, A., Inoue, K., Larom, S., Flores-uribe, J., Singh, M., Konno, M., Tomida, S., Philosof, A., Sharon, I., Yutin, N., et al. (2018). discovered using functional metagenomics.

Ran, T., Ozorowski, G., Gao, Y., Sineshchekov, O.A., Wang, W., Spudich, J.L., and Luecke, H. (2013). Cross-protomer interaction with the photoactive site in oligomeric proteorhodopsin complexes. Acta Crystallogr. D Biol. Crystallogr. 69, 1965–1980.

Renault, R., Sukenik, N., Descroix, S., Malaquin, L., Viovy, J.-L., Peyrin, J.-M., Bottani, S., Monceau, P., Moses, E., and Vignes, M. (2015). Combining Microfluidics, Optogenetics and Calcium Imaging to Study Neuronal Communication In Vitro. PLOS ONE 10, e0120680.

Ritchie, T.K., Grinkova, Y. V., Bayburt, T.H., Denisov, I.G., Zolnerciks, J.K., Atkins, W.M., and Sligar, S.G. (2009). Chapter 11 Reconstitution of Membrane Proteins in Phospholipid Bilayer Nanodiscs (Elsevier Inc.).

Robert, X., and Gouet, P. (2014). Deciphering key features in protein structures with the new ENDscript server. Nucleic Acids Res. 42, W320–W324.

Sharon, I., Alperovitch, A., Rohwer, F., Haynes, M., Glaser, F., Atamna-Ismaeel, N., Pinter, R.Y., Partensky, F., Koonin, E.V., Wolf, Y.I., et al. (2009). Photosystem I gene cassettes are present in marine virus genomes. Nature 461, 258–262.

Shcherbo, D., Merzlyak, E.M., Chepurnykh, T.V., Fradkov, A.F., Ermakova, G.V., Solovieva, E.A., Lukyanov, K.A., Bogdanova, E.A., Zaraisky, A.G., Lukyanov, S., et al. (2007). Bright far-red fluorescent protein for whole-body imaging. Nat. Methods 4, 741–746.

Shevchenko, V., Mager, T., Kovalev, K., Polovinkin, V., Alekseev, A., Juettner, J., Chizhov, I., Bamann, C., Vavourakis, C., Ghai, R., et al. (2017). Inward H+pump xenorhodopsin: Mechanism and alternative optogenetic approach. Sci. Adv. 3, 1–11.

Shigemura, S., Hososhima, S., Kandori, H., and Tsunoda, S.P. (2019). Ion Channel Properties of a Cation Channelrhodopsin, Gt_CCR4. Appl. Sci. 9, 3440.

Short, S.M. (2012). The ecology of viruses that infect eukaryotic algae. Environ. Microbiol. 14, 2253–2271.

Sineshchekov, O.A., Jung, K.-H., and Spudich, J.L. (2002). Two rhodopsins mediate phototaxis to low- and high-intensity light in Chlamydomonas reinhardtii. Proc. Natl. Acad. Sci. 99, 8689–8694.

Sineshchekov, O.A., Govorunova, E.G., and Spudich, J.L. (2009). Photosensory Functions of Channelrhodopsins in Native Algal Cells. Photochem. Photobiol. 85, 556–563.

Sineshchekov, O.A., Govorunova, E.G., Li, H., and Spudich, J.L. (2017a). Bacteriorhodopsin-like channelrhodopsins: Alternative mechanism for control of cation conductance. Proc. Natl. Acad. Sci. 114, E9512– E9519.

Sineshchekov, O.A., Govorunova, E.G., Li, H., and Spudich, J.L. (2017b). Bacteriorhodopsin-like channelrhodopsins: Alternative mechanism for control of cation conductance. Proc. Natl. Acad. Sci. 114, E9512– E9519.

Stamatakis, A.M., Schachter, M.J., Gulati, S., Zitelli, K.T., Malanowski, S., Tajik, A., Fritz, C., Trulson, M., and Otte, S.L. (2018). Simultaneous Optogenetics and Cellular Resolution Calcium Imaging During Active Behavior Using a Miniaturized Microscope. Front. Neurosci. 12, 496.

Tritsch, N.X., Granger, A.J., and Sabatini, B.L. (2016). Mechanisms and functions of GABA co-release. Nat. Rev. Neurosci. 17, 139–145.

Tsukamoto, T., Inoue, K., Kandori, H., and Sudo, Y. (2013). Thermal and Spectroscopic Characterization of a Proton Pumping Rhodopsin from an Extreme Thermophile. J. Biol. Chem. 288, 21581–21592.

Tsunogai, S., Yamahata, H., Kudo, S., and Saito, O. (1973). Calcium in the Pacific Ocean. Deep Sea Res. Oceanogr. Abstr. 20, 717–726.

Uitz, J., Claustre, H., Gentili, B., and Stramski, D. (2010). Phytoplankton class-specific primary production in the world’s oceans: Seasonal and interannual variability from satellite observations: PHYTOPLANKTON CLASS-SPECIFIC PRODUCTION. Glob. Biogeochem. Cycles 24, n/a-n/a.

Volkov, O., Kovalev, K., Polovinkin, V., Borshchevskiy, V., Bamann, C., Astashkin, R., Marin, E., Popov, A., Balandin, T., Willbold, D., et al. (2017). Structural insights into ion conduction by channelrhodopsin 2. Science 358.

Wietek, J., Wiegert, J.S., Adeishvili, N., Schneider, F., Watanabe, H., Tsunoda, S.P., Vogt, A., Elstner, M., Oertner, T.G., and Hegemann, P. (2014). Conversion of Channelrhodopsin into a Light-Gated Chloride Channel. Science 344, 409–412.

Wright, E.S. (2015). DECIPHER: Harnessing local sequence context to improve protein multiple sequence alignment. BMC Bioinformatics 16, 322.

Yau, S., Lauro, F.M., DeMaere, M.Z., Brown, M.V., Thomas, T., Raftery, M.J., Andrews-Pfannkoch, C., Lewis, M., Hoffman, J.M., Gibson, J.A., et al. (2011). Virophage control of antarctic algal host–virus dynamics. Proc. Natl. Acad. Sci. 108, 6163–6168.

Yutin, N., and Koonin, E. V (2012). Proteorhodopsin genes in giant viruses. Biol. Direct 7, 34.

